# Meta-analysis of epigenome-wide associations between DNA methylation at birth and childhood cognitive skills

**DOI:** 10.1101/2020.12.18.423421

**Authors:** Doretta Caramaschi, Alexander Neumann, Andres Cardenas, Gwen Tindula, Silvia Alemany, Lea Zillich, Giancarlo Pesce, Jari M.T. Lahti, Alexandra Havdahl, Rosa Mulder, Janine F. Felix, Henning Tiemeier, Lea Sirignano, Josef Frank, Stephanie H. Witt, Marcella Rietschel, Michael Deuschle, Karen Huen, Brenda Eskenazi, Tabea Sarah Send, Muriel Ferrer, Maria Gilles, Maria de Agostini, Nour Baïz, Sheryl L. Rifas-Shiman, Tuomas Kvist, Darina Czamara, Samuli T. Tuominen, Caroline L. Relton, Dheeraj Rai, Stephanie J. London, Katri Räikkönen, Nina Holland, Isabella Annesi-Maesano, Fabian Streit, Marie-France Hivert, Emily Oken, Jordi Sunyer, Charlotte Cecil, Gemma Sharp

**Affiliations:** Medical Research Council Integrative Epidemiology Unit (MRC IEU), Bristol Medical School, Population Health Science, University of Bristol, UK; Department of Child and Adolescent Psychiatry/Psychology, Erasmus University Medical Center Rotterdam, the Netherlands; Lady Davis Institute for Medical Research, Jewish General Hospital, Montreal, QC, Canada; Division of Environmental Health Sciences, School of Public Health, University of California, Berkeley, Berkeley, CA, USA; Children’s Environmental Health Laboratory, Division of Environmental Health Sciences, School of Public Health, University of California, Berkeley, CA, USA; I.S. Global, Barcelona Institute for Global Health, Barcelona, Spain; Universitat Pompeu Fabra (UPF), Barcelona, Spain; CIBER Epidemiology and Public Health (CIBERESP), Madrid, Spain; Department of Genetic Epidemiology in Psychiatry, Central Institute of Mental Health, Medical Faculty Mannheim, University of Heidelberg, Germany; Epidemiology of Allergic and Respiratory DIseases Team (EPAR), Institute Pierre Louis of Epidemiology and Public Health, UMR-S 1136 INSERM & Sorbonne Université, France; Department of Psychology and Logopedics, Faculty of Medicine, University of Helsinki, Finland; The Generation R Study Group, Erasmus MC, University Medical Center Rotterdam, Rotterdam, the Netherlands; Department of Pediatrics, Erasmus MC, University Medical Center Rotterdam, Rotterdam, the Netherlands; Department of Psychiatry and Psychotherapy, Central Institute of Mental Health, Medical Faculty Mannheim, University of Heidelberg, Mannheim, Germany; Center for Environmental Research and Children’s Health (CERCH), School of Public Health, University of California, Berkeley, CA, USA; Inserm, Centre for Research in Epidemiology and StatisticS (CRESS), Research team on Early life origins of health (EAROH), Villejuif, France; Division of Chronic Disease Research Across the Lifecourse, Department of Population Medicine, Harvard Medical School and Harvard Pilgrim Health Care Institute, Boston, MA, USA; Dept. Translational Research in Psychiatry, Max-Planck-Institute of Psychiatry, Munich, Germany; National Institute of Environmental Health Sciences, National Institutes of Health, Department of Health and Human Services, Research Triangle Park, NC, USA; Hospital del Mar Medical Research Institute (IMIM), Barcelona, Spain; Department of Epidemiology, Erasmus MC-University Medical Center Rotterdam, Rotterdam, the Netherlands

**Keywords:** Cognitive skills, IQ, ALSPAC, PACE, EWAS, DNA methylation, meta-analysis, cord blood

## Abstract

Cognitive skills are a strong predictor of a wide range of later life outcomes. Genetic and epigenetic associations across the genome explain some of the variation in general cognitive abilities in the general population and it is plausible that epigenetic associations might arise from prenatal environmental exposures and/or genetic variation early in life. We investigated the association between cord blood DNA methylation at birth and cognitive skills assessed in children from eight pregnancy cohorts (N=2196-3798) within the Pregnancy And Childhood Epigenetics (PACE) Consortium across overall, verbal and non-verbal cognitive scores. The associations at single CpG sites were weak for all of the cognitive domains investigated. One region near *DUSP22* on chromosome 6 was associated with non-verbal cognition in a model adjusted for maternal IQ. We conclude that there is little evidence to support the idea that cord blood DNA methylation at single CpGs can predict cognitive skills and further studies are needed to confirm regional differences.

## INTRODUCTION

The human brain starts developing prenatally in the third gestational week and its maturation extends postnatally into late adolescence and most likely adulthood. These processes occur as the result of genetic and environmental factors including the interplay between them ^1^. General cognitive ability, or intelligence, often measured as intelligent quotient (IQ), shows considerable heritability, with estimates as high as 20% in infancy, which substantially increase across the lifetime to more than 70% depending on the environment ^2^. It has been shown that cognitive skills predict important long-term outcomes such as higher educational attainment, later mortality, and better physical and mental health ^3^.

Since there is considerable environmental influence on cognitive ability in early childhood ^1^, it is plausible that environmental exposures that occur in early life, either prenatally or in early childhood, play a role in shaping children’s cognitive development. Early life environmental exposures are known to be revealed in the epigenome, as reflected in changes in DNA methylation marks on cytosine nucleotides followed by guanine (CpG) across the genome. The total heritability of DNA methylation levels from single nucleotide polymorphisms has been estimated to be on average only 19% across the genome, suggesting a strong environmental component ^4^. Moreover robust evidence indicate that maternal smoking and folate levels during pregnancy are associated with changes in the child’s epigenome at birth ^5, 6^. DNA methylation at birth can therefore reflect exposure to adverse factors, which in turn could lead to neurodevelopmental consequences.

A recent epigenome-wide association study (EWAS) of cognitive measures in older-aged adults across several cohorts, and with final sample sizes ranging between 2557 and 6809 participants, identified associations at two single CpGs ^7^. Another study of educational attainment in 10767 adults, although not specific to cognitive skills, revealed associations at nine CpG sites which are all known to be associated with smoking ^8^. At present, it is not known whether there is an association between childhood cognitive functioning and DNA methylation at birth, and whether any association found might be indicative of prenatal exposures rather than environmental exposures across the lifetime such as own smoking habits.

In this study, we aimed to investigate whether: 1) DNA methylation at the single CpG level in cord blood is prospectively associated with cognitive skills in childhood; 2) DNA methylation at the regional level in cord blood is associated with cognitive skills in childhood; 3) the association of DNA methylation in cord blood show correspondence with DNA methylation in brain.

## METHODS

### Study sample

The data used in this study were obtained from the participants of eight longitudinal birth cohorts within the Pregnancy and Childhood Epigenetics (PACE) Consortium ^9^. The cohorts were: Avon Longitudinal Study of Parents and Children (ALSPAC) ^10, 11^, Center for the Health Assessment of Mothers and Children of Salinas (CHAMACOS) ^12, 13^, Etude des Déterminants pré et post natals du développement et de la santé de l’Enfant (EDEN) ^14^, Generation R ^15^, Infancia y Medio Ambiente project (INMA) ^16^, Prediction and Prevention of Preeclampsia and Intrauterine Growth Restriction study (PREDO) ^17^, Pre-Peri- and Postnatal Stress: Epigenetic impact on Depression (POSEIDON) ^18^, and Project Viva ^19^. Ethical approval for each study was obtained by local committees and consent to use their data was obtained for all participants. Approved researchers with access to individual-level data for each cohort performed in-house analyses and shared only result files with the main analysts. Access to individual-level data is available only upon request to each cohort separately and following local procedures. For more information on each cohort, ethical approval and data access procedures please refer to the Supplementary Material.

### Epigenetic data

For each cohort, cord blood was extracted during delivery and DNA was isolated according to standard protocols. DNA was then bisulfite-treated according to standard protocols and loaded onto Infinium HumanMethylation450 BeadChip arrays (Illumina, San Diego, CA). Array images were scanned and raw methylation intensities were normalized and subjected to quality control according to cohort-specific procedures. For some cohorts, batch correction was applied before analysis (PREDO, INMA and Project Viva), while for the others batch variables were included as covariates in the analysis. Full information on the methods used within each cohort is reported in the Supplementary Methods and Materials.

### Cognitive skills

Cognitive skills were measured differently across cohorts, depending on available data at cohort-average ages ranging from 4 to 9 years (see Supplementary Material and Methods for more details). Main cognitive scores and subtests were then used to represent overall, verbal and non-verbal cognitive domains.

Overall cognitive skills were measured by:

- the Wechsler Intelligence Scale for Children, 3^rd^ edition, (WISC-III) ^20^ full-scale IQ score in ALSPAC;
- the Wechsler Intelligence Scale for Children, 4^th^ edition, (WISC-IV) ^21^ full-scale IQ score in CHAMACOS and PREDO;
- the McCarthy Scales of Children’s Abilities ^22^ general cognitive index in INMA;
- the Wide Range Assessment of Visual Motor Ability (WRAVMA) test ^23^ in Project Viva;
- the Wechsler Preschool and Primary Scale of Intelligence (WPPSI), 3^rd^ edition ^24^ fullscale IQ score in EDEN and POSEIDON.

Verbal cognitive skills were measured by:

- the Wechsler Intelligence Scale for Children, 3^rd^ edition, (WISC-III) ^20^ verbal IQ scores (derived from the verbal comprehension index and working memory index, combined) in ALSPAC;
- the Wechsler Intelligence Scale for Children, 4^th^ edition, (WISC-IV) ^21^ verbal comprehension index in CHAMACOS and PREDO;
- the McCarthy Scales of Children’s Abilities ^22^ verbal index in INMA;
- the Kaufman Brief Intelligence Test 2^nd^ edition (KBIT-II) ^25^ verbal subtest in Project Viva;
- the Wechsler Preschool and Primary Scale of Intelligence (WPPSI), 3^rd^ edition ^24^ verbal IQ score in EDEN and POSEIDON.

Non-verbal cognitive skills were measured by:

- the Wechsler Intelligence Scale for Children, 3^rd^ edition, (WISC-III) ^20^ performance IQ scores (derived from the perceptual organization index and processing speed index, combined) in ALSPAC;
- the Wechsler Intelligence Scale for Children, 4^th^ edition, (WISC-IV) ^21^ perceptual reasoning index in CHAMACOS and PREDO;
- the McCarthy Scales of Children’s Abilities ^22^ perceptual-performance index in INMA;
- the Kaufman Brief Intelligence Test 2^nd^ edition (KBIT-II) ^25^ non-verbal subtest in Project Viva;
- the Wechsler Preschool and Primary Scale of Intelligence (WPPSI), 3^rd^ edition ^24^ performance IQ score in EDEN and POSEIDON;
- the Snijders-Oomen nonverbal intelligence tests, revised (SON-R) ^26^, in Generation R.

Within each cohort, the cognitive scores were transformed into standardized z-scores. Full information on the methods used within each cohort is reported in Supplementary Methods and Materials.

### Epigenome-wide association study (EWAS)

Prior to the analyses, a data analysis plan including details of the variables, the models to use, and a sample R code was distributed to the participating cohorts. Untransformed DNA methylation beta values were used as the exposure variable. Extreme outliers (> 3 x interquartile range, either side of the 25^th^ and 75^th^ percentiles) were removed.

The effect of DNA methylation at birth on childhood cognitive skills was estimated by linear regression models (*lm*() option in R) within each cohort for each CpG site individually. The main models included the covariates: child age at cognitive testing, sex, maternal age at delivery, maternal education (cohort-specific definition), birth weight, gestational age at birth, maternal smoking status during pregnancy (any smoking compared to no smoking), parity at delivery (1 or more previous children compared to none), batch covariates (cohort-specific definition) and proportions of seven blood cell types estimated using the Houseman algorithm using a published reference dataset for cord blood ^27^.

As sensitivity analyses, three other models were run including other covariates in cohorts with the relevant data available: main model covariates + paternal education, main model covariates with maternal IQ in place of maternal education, and main model covariates + the first 10 principal components from children’s genomic data. Genomic PCs were not included in the main model as they were not available for all cohorts. Each model was run for overall, verbal and non-verbal cognitive skills. Details of the EWAS run within each cohort and specific information on the covariates used are in Supplementary Materials and Methods. A summary of the models and the corresponding sample sizes is available in Table 1.

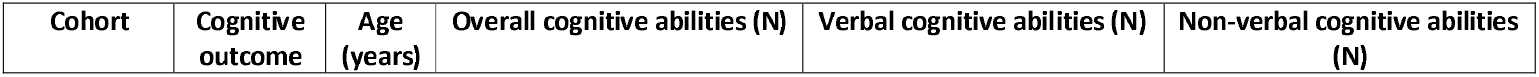

**Table 1.** Overview of the studies participating in the meta-analysis.

|  |  |  | Main | Pat.<br>Edu. | Mat.<br>IQ | PCs | Main | Pat.<br>Edu. | Mat.<br>IQ | PCs | Main | Pat.<br>Edu. | Mat.<br>IQ | PCs |
| --- | --- | --- | --- | --- | --- | --- | --- | --- | --- | --- | --- | --- | --- | --- |
| ALSPAC | WISC-III | 9 | 780 | 739 | - | 684 | 784 | 743 | - | 687 | 783 | 742 | - | 687 |
| CHAMACOS | WISC-IV | 7 | 175 | 175 | 175 | - | 175 | 175 | 175 | - | 175 | 175 | 175 | - |
| EDEN | WPPSI | 6 | 157 | 157 | - | - | 157 | 157 | - | - | 157 | 157 | - | - |
| Generation R | SON-R | 6 | - | - | - | - | - | - | - | - | 1093 | 953 | 1058 | 1058 |
| INMA | McCarthy | 4 | 319 | 319 | 311 | 280 | 319 | 319 | 311 | 280 | 319 | 319 | 311 | 280 |
| POSEIDON | WPPSI | 4 | 199 | 198 | - | 198 | 201 | 200 | - | 200 | 203 | 202 | - | 202 |
| PREDO | WISC-IV | 9 | 285 | 192 | - | 285 | 285 | 192 | - | 285 | 285 | 192 | - | 285 |
| Project VIVA | KBIT-II | 8 | 281 | 269 | 279 | - | 285 | 273 | 283 | - | 783 | 273 | 283 | - |
| **Total** |  |  | 2196 | 2049 | 765 | 1447 | 2206 | 2059 | 769 | 1452 | 3798 | 3013 | 1827 | 2512 |

### Meta-analysis

The results from the EWAS carried out in each cohort were subjected to an initial screening to check that results were comparable across cohorts (i.e. effect size distribution and precision according to sample size) using the *QCEWAS* R package ^28^. A fixed-effect metaanalysis was then performed for each model and cognitive outcome using the inverse variance weighted approach in the *metafor* R package ^29^. An independent meta-analysis was conducted at a different research institution using the *metasoft* software ^30^ to confirm the results. Heterogeneity was assessed using the *I^2^* statistics for each CpG site ^31^.

### Differential methylation region (DMR) analysis

To assess the joint effect of blood DNA methylation across different sites on cognitive skills we performed a regional analysis. The results of the meta-analysis for each model were analysed to detect differential methylation regions using the *dmrff* R package ^32^. Correlations between blood and brain methylation at CpGs within the DMRs were obtained from two online comparison tools (https://epigenetics.essex.ac.uk/bloodbrain/, date accessed 15-07-2020, and https://redgar598.shinyapps.io/BECon/, date accessed 01-12-2020). Brain expression of genes that were in DMRs was examined using the GTEx online portal (https://gtexportal.org/home/, date accessed 15-07-2020) and the Braineac online tool (http://www.braineac.org/).

## RESULTS

### Sample characteristics

All cohorts included in the main meta-analysis (Supplementary Table ST1) included a mix of male and female children (average percentage of females across cohorts ranging from 41 to 54%). Testing for childhood cognitive skills was done at a range of average ages spanning from 4 to 9 years. Maternal age at delivery was on average between 26 and 33 years overall. The children were born on average at 39-40 weeks of gestation. Maternal smoking during pregnancy varied across cohorts, with the most prevalent self-report in INMA (27%) and EDEN (25%) and the least prevalent in CHAMACOS (5%) and PREDO (4%). Maternal education was coded differently across the cohorts, with a low percentage of highly educated mothers in ALSPAC and CHAMACOS (20-21%) and highest in Generation R and Project Viva (66-78%). The percentage of children with at least one older sibling also varied across cohorts, ranging from 29 to 64%. Sample characteristics were similar for sensitivity models (Supplementary Tables ST2 and ST3). Average paternal education levels were similar to maternal ones within each cohort. Average maternal IQ was only available for 4 cohorts and it was the lowest in CHAMACOS (mean=85.7) and the highest in Project Viva (mean=111.4).

### Epigenome-wide association study meta-analysis

The results of the meta-EWAS for the main models are summarised in Figure 1 and Table 2. There was little evidence of an association between cord blood DNA methylation and childhood cognitive skills, either in terms of overall, verbal, or non-verbal scores. The most significantly associated sites did not pass the Bonferroni-adjusted p-value cut-off of 1.2 x 10^−7^. One CpG site, cg00573504 which is located in an intergenic region on chromosome 5, showed a similar association with the overall (beta = 3.71, p = 4.98E-06) and the non-verbal cognitive scores (beta = 3.38, p = 3.32E-06), whereas the other top sites (p <10^−5^) were not overlapping across the different cognitive measures. Effect sizes for the top sites were in the range of 0.02-3.6 z-score changes per 1% methylation change, whereas heterogeneity was low, with I^2^ < 60. There was some genomic inflation in the overall score model (lambda=1.11).

**Figure 1.**
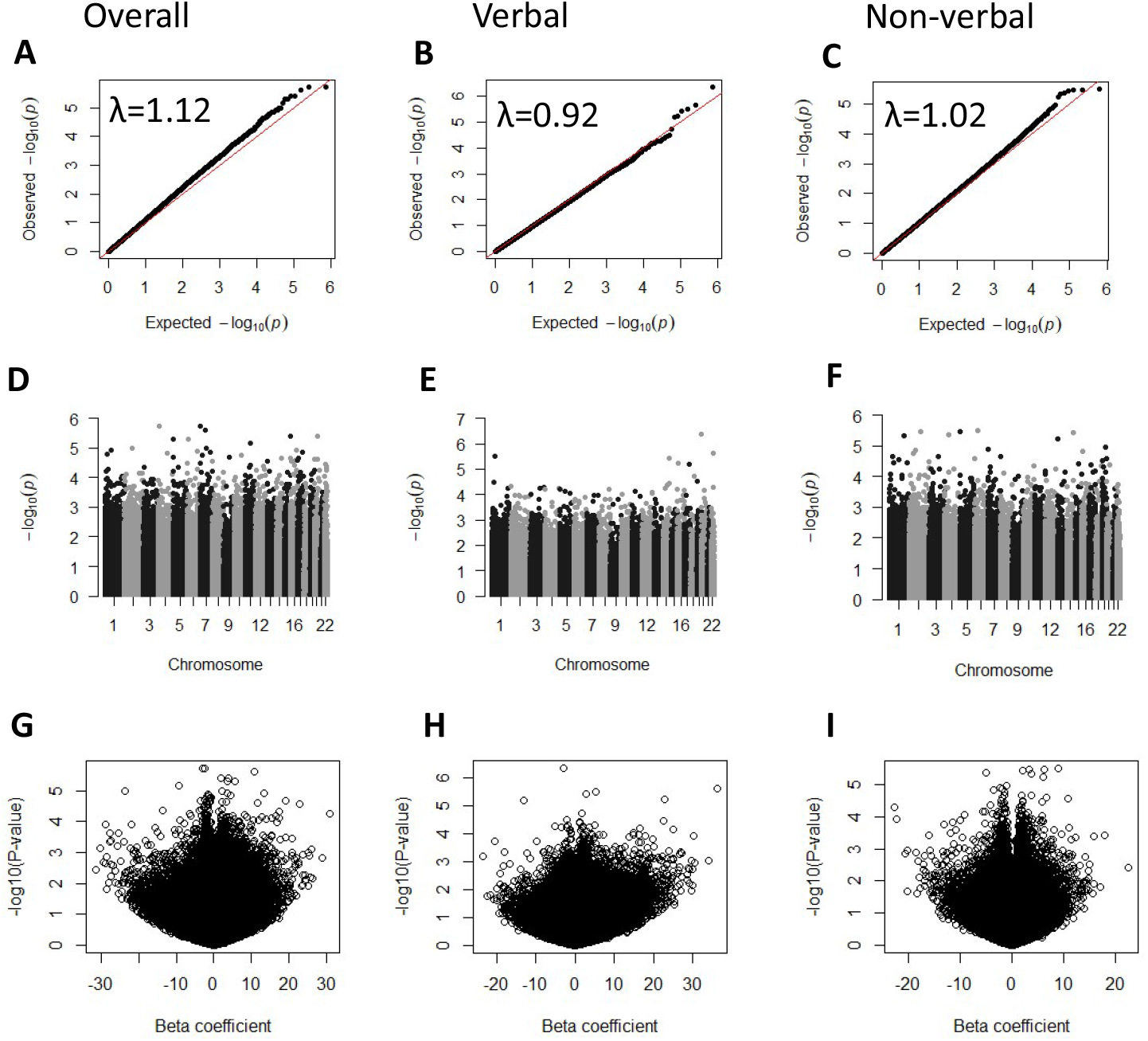
Results from the meta-analysis of epigenome-wide association studies of cognitive skills in childhood and DNA methylation in cord blood. A-C: QQ plots showing the observed vs expected probabilities per CpG site and λ index of genomic inflation. D-F: Manhattan plots showing the probability values at each CpG site according to chromosome location. G-I: Volcano plots showing the effect sizes and probability values per CpG site. Models were adjusted for age at testing, sex, maternal age at delivery, maternal education, birthweight, gestational age, maternal smoking status during pregnancy, parity, batch covariates and cell proportions.

**Table 2.** Top CpG sites (P_meta_<10^−5^) from the meta-analysis of epigenome-wide association studies of overall, verbal and non-verbal cognitive skills in childhood and DNA methylation in cord blood.

|  | CpG site | N | Beta<sup>a</sup> | S.E. | P-value<sup>b</sup> | I<sup>2</sup><sup>c</sup> | Chr. | Position | Gene<sup>d</sup> |
| --- | --- | --- | --- | --- | --- | --- | --- | --- | --- |
| **Overall** | cg05827775 | 2193 | -3.05 | 0.64 | 1.84E-06 | 0 | 4 | 9762166 |  |
|  | cg00213080 | 2190 | -2.44 | 0.51 | 1.89E-06 | 0 | 7 | 6204521 | *CYTH3* |
|  | cg26599274 | 2181 | 10.83 | 2.30 | 2.46E-06 | 15.1 | 7 | 66205733 | *RABGEF1* |
|  | cg23789148 | 2190 | 2.03 | 0.44 | 3.88E-06 | 0 | 15 | 97321146 |  |
|  | cg18622281 | 2188 | 3.96 | 0.86 | 3.91E-06 | 13.8 | 20 | 43977112 | *SDC4* |
|  | cg00573504 | 2181 | 3.71 | 0.81 | 4.98E-06 | 0 | 5 | 1123809 |  |
|  | cg09535605 | 2174 | 5.57 | 1.22 | 5.09E-06 | 5.1 | 6 | 17281327 | *RBM24* |
|  | cg21735491 | 2190 | -9.38 | 2.09 | 6.77E-06 | 0 | 11 | 66749665 |  |
|  | cg18075761 | 2152 | -23.68 | 5.36 | 9.84E-06 | 0 | 7 | 75617028 | *TMEM120A* |
| **Verbal** | cg03568675 | 2201 | -3.00 | 0.59 | 4.36E-07 | 12.3 | 20 | 979279 | *RSPO4* |
|  | cg12361663 | 2190 | 36.38 | 7.70 | 2.30E-06 | 0 | 22 | 38142561 | *TRIOBP* |
|  | cg17223866 | 2188 | 5.21 | 1.12 | 3.14E-06 | 0 | 1 | 44794158 | *ERI3* |
|  | cg11005998 | 2199 | 2.94 | 0.64 | 3.87E-06 | 0 | 14 | 101514642 | *MIR655* |
|  | cg10620273 | 2200 | 22.72 | 5.00 | 5.65E-06 | 51.0 | 16 | 3096488 | *MMP25* |
|  | cg16047144 | 2200 | -13.16 | 2.92 | 6.33E-06 | 0 | 17 | 62097953 | *ICAM2* |
| **Non-verbal** | cg04783204 | 3286 | 8.90 | 1.91 | 3.11E-06 | 53.3 | 6 | 44191600 | *SLC29A1* |
|  | cg04229103 | 3278 | 6.34 | 1.36 | 3.31E-06 | 0 | 2 | 145090268 | *GTDC1* |
|  | cg00573504 | 3282 | 3.38 | 0.73 | 3.32E-06 | 0 | 5 | 1123809 |  |
|  | cg25990848 | 3276 | 2.07 | 0.45 | 3.59E-06 | 17.8 | 14 | 105517573 | *GPR132* |
|  | cg08529049 | 3254 | -4.95 | 1.08 | 4.30E-06 | 25.6 | 4 | 48988038 | *CWH43* |
|  | cg03332597 | 3297 | 3.83 | 0.84 | 4.80E-06 | 37.3 | 1 | 185125704 | *C1orf25* |
|  | cg07489869 | 3256 | 5.94 | 1.31 | 5.85E-06 | 1.8 | 13 | 20806730 | *GJB6* |

<sup>a</sup> Beta coefficient from the regression indicating the change in IQ score standard deviations per 100% methylation change. Models were adjusted for age at testing, sex, maternal age at delivery, maternal education, birth weight, gestational age, maternal smoking status during pregnancy, parity, batch covariates and cell proportions
<sup>b</sup> Unadjusted p-value
<sup>c</sup> Heterogeneity statistics
<sup>d</sup> Gene annotation from the Illumina 450K manifest file

The results of the sensitivity models are summarised in Supplementary Figures SF1 and SF2 and in Supplementary Tables ST4, ST5, ST6 and ST7. Additionally adjusting for paternal education did not substantially change the results, with only 1 CpG showing a reduction in effect estimates of more than 1 z-score unit, whereas adjusting for maternal IQ instead of maternal education attenuated the association at 13 CpG sites and increased it at 1 site (>1 z-score unit change in effect estimates). In general, the associations, when adjusted for maternal IQ, were weaker (higher p-values). However, maternal IQ-adjusted models were more underpowered due to a smaller sample size. The associations with nonverbal scores were less affected by the adjustment by maternal IQ (3 sites at p-value<0.001). Adjusting for child’s genomic PCs reduced inflation (lambdas 0.94-1.05) and revealed stronger effects at 2 sites (>1 z-score unit increase).

Using the results of the main models, we looked up findings from recent EWAS metaanalyses of adult blood cells methylation and either cognitive skills or educational attainment, both in adulthood. The two main CpG sites from the cognitive skills EWAS did not replicate in our study (p-value>0.05). We were able to replicate the previously identified association of DNA methylation at cg17759224 (intergenic region on chromosome 1) with non-verbal scores (p=0.00065), out of the 34 sites found across the various scores in the original study (Supplementary Table ST8). None of the CpG sites from the educational attainment EWAS had cord blood DNA methylation levels associated with childhood cognitive outcomes in our meta-analysis (Supplementary Table ST9).

### DMR analysis

We performed a DMR analysis to investigate the association of DNA methylation at clusters of CpG sites with cognitive skills and found little evidence of clusters in the main models across the cognitive scores. When we adjusted for maternal IQ (3 cohorts for overall and verbal cognitive skills, 4 cohorts for non-verbal cognitive skills), methylation within a region comprising 5 CpG sites on chromosome 6 was associated with slightly higher non-verbal scores (Supplementary Table ST10). The heterogeneity at these sites was very low (*I^2^* = 0-5). This region is located within the *DUSP22* gene, a phosphatase that is expressed across examined tissues including blood and several brain areas (Supplementary Figure SF3). DNA methylation in blood within this region highly correlates with brain methylation at all CpGs across all brain regions and across the two datasets interrogated, with r >0.9, p < 10^−32^ in the prefrontal cortex, entorhinal cortex, cerebellum and superior temporal gyrus (Supplementary Table ST11), and r values of 0.4-0.8, reaching the 90^th^ percentile, for BA10, BA20 and BA7 (Supplementary Figure SF4). A search within two mQTL databases (www.mqtldb.org and mqtldb.godmc.org.uk/) showed only *trans*-mQTLs for these CpG sites, therefore we could not perform any Mendelian randomization analyses to investigate if these associations are causal.

## DISCUSSION

We have performed the largest epigenome-wide association study of cognitive skills in cord blood, by running individual EWAS in eight cohorts and combining the results through meta-analysis. We hypothesized that, at birth, we would be able to identify methylation marks that predict variation in cognitive skills in childhood. Overall, the evidence at single CpG sites was weak across all models to confirm an association. In one region spanning 5 CpGs on chromosome 6 methylation was correlated with small increases in non-verbal scores after adjusting for maternal IQ. This association was revealed in a subgroup sensitivity analysis, with limited power compared to the main analyses (3-4 cohorts instead of 7-8). Cord blood DNA methylation in this region was highly correlated with brain methylation, although we could not identify cis-mQTLs to perform further analyses to establish if methylation in this region is causal to variation in cognitive skills.

The differentially methylated region on chromosome 6 is located in the *DUSP22* gene, coding for a phosphatase expressed ubiquitously, including in blood and across brain regions. From previously published data ^33^, there is evidence that, in whole blood, DNA methylation within this region is associated with increased *DUSP22* expression (Supplementary Table ST12). We also observed that DNA methylation correlates highly between peripheral blood and brain tissue. Moreover, methylation at the *DUSP22* gene in brain tissue has been previously implicated in schizophrenia, Parkinson’s and Alzheimer’s disease, albeit in different directions depending on the brain region investigated ^34–36^. These findings, altogether, suggest that DNA methylation may affect brain functioning via changes in *DUSP22* expression. However, in our study this association was observed in only one of the sensitivity models, with a smaller sample size than the other models since not all the cohorts had data available on maternal IQ. It has been previously shown that maternal IQ affects children cognitive development independently from socioeconomic status ^37^. This suggests that maternal IQ could affect the association of children cognition with methylation at birth independently from maternal education (as a proxy for socioeconomic status).

Our study replicated the association of 1 of the 34 CpG sites identified in a previous EWAS of cognitive skills in adulthood ^7^, and none of the sites found in an EWAS of educational attainment ^8^. In those previous studies, both carried out in adulthood, blood cells methylation patterns seemed to be an indication of lifestyle characteristics such as smoking and BMI. Since our study looked for methylation patterns at birth, we would not necessarily expect to see the effect of direct exposures that would occur later in life. Our study suggests that the relationship between cognitive skills and blood DNA methylation seen previously are reflective of exposures after birth, rather than the prenatal period. Another explanation is that blood methylation marks arise from a gene-environment interaction and they appear only later in life, due to the cumulative effect of environmental exposures that are moderated by genetic variation.

It is also possible that the effect of prenatal exposures on childhood cognitive skills are associated with brain DNA methylation patterns that are not captured by cord blood DNA methylation. Largely due to the inaccessibility of brain tissue, most molecular studies of brain-related traits and disorders rely on blood samples, including cord blood in studies of children at birth. Although DNA methylation patterns are often tissue-specific ^38^, there are strong cross-tissue correlations at specific sites, and studies on blood-brain correspondence allow us to make comparisons that are relevant to brain phenotypes ^39–41^. Moreover, despite blood not being the main target tissue for neurodevelopmental conditions, epigenetic associations have been found in peripheral blood e.g. for schizophrenia and autism ^42, 43^ and correspondence with brain is not a prerequisite for functionally-relevant DNA methylation changes where immune-related effects on the brain are potential mechanisms Additionally, peripheral tissue can still serve as a biomarker, when stable signals are identified.

The main strength of this study is the large sample size achieved by analysing data from eight cohorts and combining the results in a meta-analysis allowing to identify only robust results. By using the same protocol and script in all the cohorts we have reduced bias due to heterogeneity.

Our study also has some limitations. Cognitive skills were assessed using different instruments across the cohorts investigated, which can explain some heterogeneity and reduced precision in our estimates. Although the sample size was large, some cohorts could not participate in some of the models, such as Generation R only having data on non-verbal skills and ALSPAC not having maternal IQ data. Our study consisted mostly of participants that identified themselves as white and lived either in Europe or the US. Therefore, the results of this study are not generalizable globally to other ethnic groups and countries. Moreover, the 450K methylation arrays only capture 2% of the CpGs in the human genome.

In conclusion, we have conducted the largest epigenome-wide scan at birth for cognitive functioning in childhood. We found evidence for a regional association of DNA methylation at birth with childhood non-verbal cognitive skills. Overall, the evidence does not suggest that cord blood DNA methylation at the single CpGs investigated could be an indication of later cognitive skills, either overall, verbal or non-verbal. Most likely, other DNA methylation marks associated with cognition in peripheral blood arise later in life or are stochastic. Further studies are needed to replicate these results across more ethnically diverse cohorts, in larger samples with data on maternal IQ and using higher resolution arrays.

## ACKNOWLEDGEMENTS

We are extremely grateful to all the families who took part in the study, the midwives for their help in recruiting them, interviewers, computer and laboratory technicians, clerical workers, research scientists, volunteers, managers, receptionists and nurses, general practitioners, hospitals and pharmacies. In particular, we thank: Dr S.M. Ring, Dr W. McArdle, Dr M. Suderman (ALSPAC); Mr. Michael Verbiest, Ms. Mila Jhamai, Ms. Sarah Higgins, Mr. Marijn Verkerk, Dr. Lisette Stolk and Dr. A.Teumer (Generation R); Silvia Fochs and Nuria Pey (INMA); L. Douhaud, S. Bedel, B. Lortholary, S. Gabriel, M. Rogeon, M. Malinbaum. J.Y Bernard, J. Botton, M.A. Charles, P. Dargent-Molina, B. de Lauzon-Guillain, P. Ducimetière, B. Foliguet, A. Forhan, X. Fritel, A. Germa, V. Goua, R. Hankard, B. Heude, M. Kaminski, B. Larroque†, N. Lelong, J. Lepeule, G. Magnin, L. Marchand, C. Nabet, F. Pierre, R. Slama, M.J. Saurel-Cubizolles, M. Schweitzer, O. Thiebaugeorges (EDEN); E. Hamäläinen, E. Kajantie, H. Laivuori, P.M. Villa, A-K. Pesonen, A. Aitokallio-Tallberg, A-M. Henry, V.K. Hiilesmaa, T. Karipohja, R. Meri, S. Sainio, T. Saisto, S. Suomalainen-Konig, V-M. Ulander, T. Vaitilo L. Keski-Nisula, M-R. Orden, E. Koistinen, T. Walle, R. Solja, M. Kurkinen, P. Taipale. P. Staven and J. Uotila (PREDO). Please see the Supplementary Material for more detailed acknowledgements.

## DATA AVAILABILITY

Meta-analysis results files will be deposited in the EWAS Catalog data repository upon publication. Individual-level data are available upon request to the cohorts involved and according to their procedures.

## AUTHOR CONTRIBUTION

D.Ca. study design, ALSPAC analysis, meta-analysis, interpretation of results, manuscript drafting and revision; A.N. shadow meta-analysis, Generation R analysis, interpretation of results, critical revision of manuscript; A.C. Project Viva analysis and interpretation; G.T. CHAMACOS analysis, manuscript review; S.A. INMA analysis, manuscript revision; L.Z. POSEIDON analysis, manuscript revision; G.P. EDEN analysis, manuscript revision; J.M.T.L provided funding, PREDO data acquisition, quality control, cohort description, reviewed manuscript; A.H. contributed to analysis plan and revision of drafts and manuscript; R.M. Generation R data acquisition, data analysis, critical revision of the manuscript; J.F.F. Interpretation of results, critical revision of manuscript; H.T. Generation R data acquisition, reviewing drafts, provided funding, quality control and supervision of IQ data; L.S. data acquisition, manuscript revision; J.F. POSEIDON data acquisition, manuscript revision; S.H.W. POSEIDON data acquisition, manuscript revision; M.R. POSEIDON data acquisition, manuscript revision; M.D. POSEIDON data acquisition, manuscript revision; K. H. POSEIDON manuscript revision; B.E. manuscript revision; T.S.S. POSEIDON data acquisition, manuscript revision; M.F. INMA data collection, manuscript revision; M.G. POSEIDON data acquisition, manuscript revision; M.deA. EDEN cohort data collection; N.B. EDEN cohort data analysis; S.L.R-F Project Viva analysis and interpretation; T.K. data acquisition, PREDO data collection, reviewed manuscript; D.Cz. PREDO data acquisition and cleaning, manuscript review; C.L.R. ALSPAC study funding and data acquisition, interpretation of findings; D.R. study design and analysis plan; S.T.T. PREDO data analyses, reviewed manuscript; S.J.L. interpretation and manuscript review; N.H. manuscript review; K.R. provided funding, planned the PREDO data collection, reviewed manuscript; I.A-M EDEN cohort data collection and funding acquisition; F.S. manuscript review; M-F.H. Project Viva analysis and interpretation; E.O. Project Viva analysis and interpretation; J.S. funding, manuscript revision; C.C. study design, manuscript drafting; G.S. study design, manuscript drafting.

