## supplementary tables for "Meta-analysis of epigenome-wide associations between DNA methylation at birth and childhood cognitive skills"

**Supplementary Table ST1**. Sample characteristics across the cohorts used in the meta-analysis EWAS for cognitive abilities (Main models: O= Overall, V=Verbal, NV= Non-Verbal).

|  | **ALSPAC** | | | **CHAMACOS** | | | **EDEN** | | | **Gen. R** | **INMA** | | | **POSEIDON** | | | **PREDO** | | | **Project VIVA** | | |
| --- | --- | --- | --- | --- | --- | --- | --- | --- | --- | --- | --- | --- | --- | --- | --- | --- | --- | --- | --- | --- | --- | --- |
|  | **O** | **V** | **NV** | **O** | **V** | **NV** | **O** | **V** | **NV** | **NV** | **O** | **V** | **NV** | **O** | **V** | **NV** | **O** | **V** | **NV** | **O** | **V** | **NV** |
| **N** | 780 | 784 | 783 | 175 | 175 | 175 | 157 | 157 | 157 | 1093 | 319 | 319 | 319 | 199 | 201 | 203 | 285 | 285 | 285 | 281 | 285 | 285 |
| **Sex (% females)** | 52 | 52 | 51 | 54 | 54 | 54 | 41 | 41 | 41 | 51 | 48 | 48 | 48 | 56 | 56 | 56 | 49 | 49 | 49 | 50 | 50 | 50 |
| **Age at test, mean (SD)** | 8.6  (0.2) | 8.6  (0.2) | 8.6  (0.2) | 7.1  (0.2) | 7.1  (0.2) | 7.1  (0.2) | 5.7  (0.1) | 5.7  (0.1) | 5.7  (0.1) | 6.1  (0.3) | 4.5  (0.2) | 4.5  (0.2) | 4.5  (0.2) | 3.7  (0.1) | 3.7  (0.1) | 3.7  (0.1) | 8.7  (0.8) | 8.7  (0.8) | 8.7  (0.8) | 7.8  (0.7) | 7.8  (0.8) | 7.8  (0.7) |
| **Age at delivery, mean (SD)** | 29.8  (4.4) | 29.8  (4.4) | 29.8  (4.4) | 26.4  (5.3) | 26.4  (5.3) | 26.4  (5.3) | 30.3  (5.0) | 30.3  (5.0) | 30.3  (5.0) | 31.9  (4.1) | 30.4  (4.1) | 30.4  (4.1) | 30.4  (4.1) | 32.0  (5.0) | 32.0  (4.6) | 31.7  (4.6) | 34.0  (5.5) | 34.0  (5.5) | 34.0  (5.5) | 33.3  (4.4) | 33.3  (4.3) | 33.3  (4.3) |
| **Gestational age, mean (SD)** | 39.6  (1.5) | 39.6  (1.5) | 39.6  (1.5) | 39 .0  (1.4) | 39.0  (1.4) | 39.0  (1.4) | 39.5  (1.4) | 39.5  (1.4) | 39.5  (1.4) | 40.2  (1.5) | 39.8  (1.3) | 39.8  (1.3) | 39.8  (1.3) | 39.2  (1.1) | 39.2  (1.2) | 39.2  (1.2) | 39.8  (1.6) | 39.7  (1.6) | 39.7  (1.6) | 39.9  (1.5) | 39.9  (1.5) | 39.9  (1.5) |
| **Smoking in pregnancy (%)** | 13 | 13 | 13 | 5 | 5 | 5 | 25 | 25 | 25 | 20 | 27 | 27 | 27 | 24 | 23 | 24 | 4 | 4 | 4 | 7 | 7 | 7 |
| **Maternal education**  **(N per group or mean and SD)*** | 619 | 622 | 622 | 137 | 137 | 137 | 38 | 38 | 38 | 12 | 78 | 78 | 78 | 16.8  (3.5) | 16.8  (3.5) | 16.8  (3.5) | 0 | 0 | 0 | 62 | 63 | 63 |
|  | 161 | 162 | 161 | 38 | 38 | 38 | 67 | 67 | 67 | 89 | 137 | 137 | 137 |  |  |  | 3 | 3 | 3 | 219 | 222 | 222 |
|  |  |  |  |  |  |  | 52 | 52 | 52 | 260 | 104 | 104 | 104 |  |  |  | 103 | 103 | 103 |  |  |  |
|  |  |  |  |  |  |  |  |  |  | 297 |  |  |  |  |  |  | 72 | 72 | 72 |  |  |  |
|  |  |  |  |  |  |  |  |  |  | 435 |  |  |  |  |  |  | 107 | 107 | 107 |  |  |  |
|  |  |  |  |  |  |  |  |  |  |  |  |  |  |  |  |  | 0 | 0 | 0 |  |  |  |
| **Parity (% at least one sibling)** | 53 | 53 | 53 | 64 | 64 | 64 | 58 | 58 | 58 | 38 | 41 | 41 | 41 | 49 | 49 | 49 | 29 | 29 | 29 | 48 | 48 | 48 |

* ALSPAC: low= no university degree, high= university degree; CHAMACOS: low= less than high school, high=at least high school; EDEN: low= upper secondary education, middle= post-secondary non-tertiary, high=tertiary; Generation R= continuous education level from low to high (primary, secondary-phase 1, secondary-phase 2, higher-phase 1, higher-phase 2); INMA: low, middle, high; POSEIDON: years of education (mean, SD); PREDO: groups 1-6 (7-9 had N=0) from high to low education (continuous); Project VIVA: low= not college graduate, high= college graduate.

**Supplementary Table ST2**. Sample characteristics across the cohorts used in the meta-analysis EWAS for cognitive abilities (Sensitivity models further adjusted for paternal education: O= Overall, V=Verbal, NV= Non-Verbal).

|  | **ALSPAC** | | | **CHAMACOS** | | | **EDEN** | | | **Gen. R** | **INMA** | | | **POSEIDON** | | | **PREDO** | | | **Project VIVA** | | |
| --- | --- | --- | --- | --- | --- | --- | --- | --- | --- | --- | --- | --- | --- | --- | --- | --- | --- | --- | --- | --- | --- | --- |
|  | **O** | **V** | **NV** | **O** | **V** | **NV** | **O** | **V** | **NV** | **NV** | **O** | **V** | **NV** | **O** | **V** | **NV** | **O** | **V** | **NV** | **O** | **V** | **NV** |
| **N** | 739 | 743 | 742 | 175 | 175 | 175 | 157 | 157 | 157 | 953 | 319 | 319 | 319 | 198 | 200 | 202 | 192 | 192 | 192 | 269 | 273 | 273 |
| **Sex (% females)** | 51 | 51 | 51 | 54 | 54 | 54 | 41 | 41 | 41 | 52 | 48 | 48 | 48 | 57 | 56 | 56 | 50 | 50 | 50 | 51% | 51% | 51% |
| **Age at test, mean (SD)** | 8.6  (0.2) | 8.6  (0.2) | 8.6  (0.2) | 7.1  (0.2) | 7.1  (0.2) | 7.1  (0.2) | 5.7  (0.1) | 5.7  (0.1) | 5.7  (0.1) | 6.1  (0.3) | 4.5  (0.2) | 4.5  (0.2) | 4.5  (0.2) | 44.9  (0.9) | 44.9  (0.9) | 44.9  (0.9) | 8.5  (0.7) | 8.5  (0.7) | 8.5  (0.7) | 7.8 (0.7) | 7.8 (0.7) | 7.8 (0.7) |
| **Age at delivery, mean (SD)** | 29.8  (4.3) | 29.8  (4.4) | 29.9  (4.4) | 26.4  (5.3) | 26.4  (5.3) | 26.4  (5.3) | 30.3  (5.0) | 30.3  (5.0) | 30.3  (5.0) | 31.8  (4.0) | 30.4  (4.1) | 30.4  (4.1) | 30.4  (4.1) | 31.8  (4.5) | 31.8  (4.5) | 31.8  (4.6) | 33.8  (5.1) | 33.8  (5.1) | 33.8  (5.1) | 33.4 (4.3) | 33.4 (4.3) | 33.4 (4.3) |
| **Gestational age, mean (SD)** | 40  (1.5) | 39.6  (1.5) | 39.6  (1.5) | 39  (1.4) | 39  (1.4) | 39  (1.4) | 39.5  (1.4) | 39.5  (1.4) | 39.5  (1.4) | 40.2  (1.5) | 39.8  (1.3) | 39.8  (1.3) | 39.8  (1.3) | 39.2  (1.2) | 39.2  (1.2) | 39.2  (1.2) | 39.7  (1.7) | 39.7  (1.7) | 39.7  (1.7) | 40.0 (1.5) | 40.0 (1.5) | 40.0 (1.5) |
| **% any smoking in pregnancy** | 12 | 12 | 12 | 5 | 5 | 5 | 25 | 25 | 25 | 20 | 27 | 27 | 27 | 46 | 46 | 47 | 4 | 4 | 4 | 7 | 7 | 7 |
| **Maternal education**  **(N per group or mean and SD)*** | 579 | 582 | 582 | 137 | 137 | 137 | 38 | 38 | 38 | 0 | 78 | 78 | 78 | 16.8  (3.5) | 16.8  (3.5) | 16.8 (3.5) | 0 | 0 | 0 | 58 | 59 | 59 |
|  | 160 | 161 | 160 | 38 | 38 | 38 | 67 | 67 | 67 | 10 | 137 | 137 | 137 |  |  |  | 3 | 3 | 3 | 211 | 214 | 214 |
|  |  |  |  |  |  |  |  |  |  | 73 | 104 | 104 | 104 |  |  |  | 57 | 57 | 57 |  |  |  |
|  |  |  |  |  |  |  |  |  |  | 222 |  |  |  |  |  |  | 51 | 51 | 51 |  |  |  |
|  |  |  |  |  |  |  |  |  |  | 262 |  |  |  |  |  |  | 81 | 81 | 81 |  |  |  |
|  |  |  |  |  |  |  |  |  |  | 386 |  |  |  |  |  |  | 0 | 0 | 0 |  |  |  |
| **Paternal education**  **(N per group or mean and SD)*** | 532 | 536 | 535 | 148 | 148 | 148 | 48 | 48 | 48 | 0 | 111 | 111 | 111 | 16.39 (3.78) | 16.39 (3.78) | 16.39 (3.77) | 0 | 0 | 0 | 72 | 74 | 74 |
|  | 207 | 207 | 207 | 127 | 127 | 127 | 72 | 72 | 72 | 22 | 143 | 143 | 143 |  |  |  | 0 | 0 | 0 | 197 | 199 | 199 |
|  |  |  |  |  |  |  | 37 | 37 | 37 | 100 | 65 | 65 | 65 |  |  |  | 3 | 3 | 3 |  |  |  |
|  |  |  |  |  |  |  |  |  |  | 219 |  |  |  |  |  |  | 22 | 22 | 22 |  |  |  |
|  |  |  |  |  |  |  |  |  |  | 195 |  |  |  |  |  |  | 40 | 40 | 40 |  |  |  |
|  |  |  |  |  |  |  |  |  |  | 417 |  |  |  |  |  |  | 34 | 34 | 34 |  |  |  |
|  |  |  |  |  |  |  |  |  |  |  |  |  |  |  |  |  | 38 | 38 | 38 |  |  |  |
|  |  |  |  |  |  |  |  |  |  |  |  |  |  |  |  |  | 55 | 55 | 55 |  |  |  |
|  |  |  |  |  |  |  |  |  |  |  |  |  |  |  |  |  | 0 | 0 | 0 |  |  |  |
| **Parity (% at least one sibling)** | 54 | 54 | 54 | 38 | 38 | 38 | 58 | 58 | 58 | 37 | 41 | 41 | 41 | 49 | 49 | 49 | 33 | 33 | 33 | 47 | 48 | 48 |

* ALSPAC: low= no university degree, high= university degree; CHAMACOS: low= less than high school, high=at least high school; EDEN: low= upper secondary education, middle= post-secondary non-tertiary, high=tertiary; Generation R= continuous education level from low to high (primary, secondary-phase 1, secondary-phase 2, higher-phase 1, higher-phase 2); INMA: low, middle, high; POSEIDON: years of education (mean, SD); PREDO: groups 1-9 (7-9 for maternal had N=0) from high to low education (continuous); Project VIVA: low= not college graduate, high= college graduate.

**Supplementary Table ST3**. Sample characteristics across the cohorts used in the meta-analysis EWAS for cognitive abilities (Sensitivity models adjusted for maternal IQ instead of maternal education: O= Overall, V=Verbal, NV= Non-Verbal).

|  | **CHAMACOS** | | | **Gen. R** | **INMA** | | | **Project VIVA** | | |
| --- | --- | --- | --- | --- | --- | --- | --- | --- | --- | --- |
|  | **O** | **V** | **NV** | **NV** | **O** | **V** | **NV** | **O** | **V** | **NV** |
| **N** | 175 | 175 | 175 | 1058 | 311 | 311 | 311 | 279 | 283 | 283 |
| **Sex (% females)** | 54 | 54 | 54 | 51 | 49 | 49 | 49 | 50 | 50 | 50 |
| **Age at test, mean (SD)** | 7.1  (0.2) | 7.1  (0.2) | 7.1  (0.2) | 6.1  (0.3) | 4.5  (0.2) | 4.5  (0.2) | 4.5  (0.2) | 7.9  (0.7) | 7.9  (0.8) | 7.9  (0.8) |
| **Age at delivery, mean (SD)** | 26.4  (5.3) | 26.4  (5.3) | 26.4  (5.3) | 31.9  (4.1) | 30.4  (4.1) | 30.4  (4.1) | 30.4  (4.1) | 33.4  (4.4) | 33.4  (4.4) | 33.4  (4.4) |
| **Gestational age, mean (SD)** | 39  (1.4) | 39  (1.4) | 39  (1.4) | 40.2  (1.5) | 39.8  (1.3) | 39.8  (1.3) | 39.8  (1.3) | 40.0  (1.5) | 40.0  (1.5) | 40.0  (1.5) |
| **% any smoking in pregnancy** | 5 | 5 | 5 | 20 | 27 | 27 | 27 | 7 | 7 | 7 |
| **Maternal IQ score *, mean (SD)** | 85.7  (20.5) | 85.7  (20.5) | 85.7  (20.5) | 102.8  (12.1) | 10.6  (2.8) | 10.6  (2.8) | 10.6  (2.8) | 111.4  (12.3) | 111.4  (12.3) | 111.4  (12.3) |
| **Parity (% at least one sibling)** | 64 | 64 | 64 | 38 | 42 | 42 | 42 | 47 | 48 | 48 |

CHAMACOS= Peabody Vocabulary Test, Generation R= Raven’s Advanced Progressive Matrices Test, INMA= Similarities subtest from the Wechsler Adult Intelligence-3rd edition, Project VIVA= Kaufman Brief Intelligence Test 2nd edition

**Supplementary Table ST4**. Top CpG sites (p-value <10^-5^) from the meta-analysis of epigenome-wide association studies of overall, verbal and non-verbal cognitive abilities in childhood and DNA methylation in cord blood with further adjustment for paternal education.

|  | **CpG site** | **N** | **Beta**^a^ | **S.E.** | **P-value** ^b^ | **I^2^** ^c^ | **Chr.** | **Position** | **Gene** ^d^ |
| --- | --- | --- | --- | --- | --- | --- | --- | --- | --- |
| **Overall** | cg00213080 | 2043 | -2.48 | 0.52 | 1.48E-06 | 0 | 7 | 6204521 | *CYTH3* |
|  | cg23789148 | 2043 | 2.12 | 0.46 | 4.60E-06 | 0 | 15 | 97321146 |  |
|  | cg03931865 | 2043 | -1.64 | 0.36 | 5.28E-06 | 41.1 | 16 | 47048107 |  |
|  | cg23336139 | 2038 | -5.18 | 1.14 | 5.78E-06 | 46.3 | 12 | 1.18E+08 | *TESC* |
|  | cg14848685 | 2043 | -1.69 | 0.38 | 6.71E-06 | 0 | 6 | 1.39E+08 | *KIAA1244* |
|  | cg03493774 | 2049 | -1.11 | 0.25 | 6.74E-06 | 0 | 14 | 92879474 | *SLC24A4* |
|  | cg26599274 | 2034 | 10.58 | 2.36 | 7.17E-06 | 0 | 7 | 66205733 | *RABGEF1* |
|  | cg05827775 | 2046 | -2.95 | 0.66 | 7.73E-06 | 17.1 | 4 | 9762166 |  |
|  | cg18622281 | 2041 | 3.90 | 0.87 | 7.76E-06 | 5.8 | 20 | 43977112 | *SDC4* |
|  | cg13775913 | 2038 | 20.11 | 4.50 | 8.13E-06 | 0 | 17 | 7297869 | *PLSCR3* |
|  | cg21735491 | 2043 | -9.20 | 2.08 | 9.38E-06 | 0 | 11 | 66749665 |  |
| **Verbal** | cg09178369 | 2045 | 4.83 | 1.02 | 2.20E-06 | 39.9 | 6 | 7828165 | *BMP6* |
|  | cg12361663 | 2043 | 35.71 | 7.67 | 3.22E-06 | 0 | 22 | 38142561 | *TRIOBP* |
|  | cg05083414 | 2037 | -4.17 | 0.92 | 6.32E-06 | 0 | 4 | 2627039 | *FAM193A* |
|  | cg10620273 | 2053 | 22.04 | 4.97 | 9.19E-06 | 55.8 | 16 | 3096488 | *MMP25* |
|  | cg03568675 | 2054 | -2.85 | 0.64 | 9.28E-06 | 2.0 | 20 | 979279 | *RSPO4* |
|  | cg12961010 | 2049 | 3.58 | 0.81 | 9.50E-06 | 0 | 12 | 132938327 |  |
| **Non-verbal** | cg03332597 | 3011 | 4.30 | 0.86 | 6.49E-07 | 31.0 | 1 | 185125704 | *C1orf25* |
|  | cg04783204 | 3001 | 9.76 | 1.99 | 8.97E-07 | 56.3 | 6 | 44191600 | *SLC29A1* |
|  | cg25990848 | 2991 | 2.18 | 0.46 | 2.45E-06 | 0 | 14 | 105517573 | *GPR132* |
|  | cg21885231 | 3013 | 2.56 | 0.55 | 3.06E-06 | 0 | 7 | 1453743 |  |
|  | cg04229103 | 2991 | 6.58 | 1.42 | 3.74E-06 | 0 | 2 | 145090268 | *GTDC1* |
|  | cg14255471 | 2999 | -1.45 | 0.32 | 4.46E-06 | 42.6 | 14 | 185125704 |  |
|  | cg07216133 | 3003 | -2.10 | 0.46 | 4.80E-06 | 0 | 6 | 44191600 | *RREB1* |
|  | cg16355591 | 2980 | -3.99 | 0.87 | 4.92E-06 | 0 | 16 | 105517573 |  |
|  | cg17935281 | 2967 | 4.21 | 0.93 | 5.84E-06 | 30 | 3 | 1453743 | *SEMA5B* |
|  | cg00573504 | 2995 | 3.39 | 0.76 | 7.53E-06 | 0 | 5 | 145090268 |  |
|  | cg21950196 | 3013 | 2.22 | 0.50 | 9.79E-06 | 0 | 1 | 106660938 | *GUK1* |

^a^ Beta coefficient from the regression indicating the change in IQ score standard deviations per 100% methylation change. Models were adjusted also for age at testing, sex, maternal age at delivery, maternal education, birthweight, gestational age, maternal smoking status during pregnancy, parity, batch covariates and cell proportions

^b^ Unadjusted p-value

^c^ Heterogeneity statistics

^d^ Gene annotation from the Illumina 450K manifest file

**Supplementary Table ST5**. Top CpG sites (p-value <10^-5^) from the meta-analysis of epigenome-wide association studies of overall, verbal and non-verbal cognitive abilities in childhood and DNA methylation in cord blood with adjustment for maternal IQ.

|  | **CpG site** | **N** | **Beta**^a^ | **S.E.** | **P-value** ^b^ | **I^2^** ^c^ | **Chr.** | **Position** | **Gene** ^d^ |
| --- | --- | --- | --- | --- | --- | --- | --- | --- | --- |
| **Overall** | - | - | - | - | - | - | - | - | *-* |
| **Verbal** | cg16713732 | 769 | 3.85 | 0.84 | 4.32E-06 | 18.0 | 3 | 127325013 | *MCM2* |
| **Non-verbal** | cg10254690 | 1548 | -5.68 | 1.27 | 8.08E-06 | 43.3 | 10 | 126107861 | *OAT* |
|  | cg10273821 | 1579 | 2.19 | 0.49 | 8.97E-06 | 0 | 15 | 28014188 | *OCA2* |

^a^ Beta coefficient from the regression indicating the change in IQ score standard deviations per 100% methylation change. Models were adjusted also for age at testing, sex, maternal age at delivery, birthweight, gestational age, maternal smoking status during pregnancy, parity, batch covariates and cell proportions

^b^ Unadjusted p-value

^c^ Heterogeneity statistics

^d^ Gene annotation from the Illumina 450K manifest file

**Supplementary Table ST6**. Top CpG sites (p-value <10^-5^) from the meta-analysis of epigenome-wide association studies of overall, verbal and non-verbal cognitive abilities in childhood and DNA methylation in cord blood with further adjustment for 10 principal components from genetic data.

|  | **CpG site** | **N** | **Beta**^a^ | **S.E.** | **P-value** ^b^ | **I^2^** ^c^ | **Chr.** | **Position** | **Gene** ^d^ |
| --- | --- | --- | --- | --- | --- | --- | --- | --- | --- |
| **Overall** | cg06760279 | 1442 | 3.51 | 0.75 | 3.05E-06 | 3.7 | 14 | 23947073 | *NGDN* |
|  | cg21899374 | 1442 | 3.16 | 0.68 | 3.84E-06 | 0 | 8 | 144915517 |  |
|  | cg23951474 | 1447 | -1.94 | 0.42 | 3.99E-06 | 20.6 | 11 | 2188061 | *TH* |
|  | cg23789148 | 1444 | 2.34 | 0.51 | 4.59E-06 | 0 | 15 | 97321146 |  |
|  | cg05264908 | 1425 | 2.40 | 0.53 | 5.80E-06 | 0 | 16 | 2049630 | *ZNF598* |
|  | cg13783152 | 1447 | 4.62 | 1.04 | 8.60E-06 | 0 | 9 | 17579026 | *SH3GL2* |
| **Verbal** | cg10334750 | 1432 | 4.04 | 0.86 | 2.54E-06 | 0 | 8 | 101348456 |  |
|  | cg05083414 | 1435 | -4.53 | 0.98 | 3.75E-06 | 51.0 | 4 | 2627039 | *FAM193A* |
|  | cg23951474 | 1452 | -1.95 | 0.42 | 4.29E-06 | 38.4 | 11 | 2188061 | *TH* |
|  | cg12162201 | 1451 | 38.17 | 8.56 | 8.21E-06 | 62.6 | 21 | 48055631 | *PRMT2* |
| **Non-verbal** | cg07805967 | 2510 | 3.55 | 0.74 | 1.42E-06 | 0 | 10 | 53459337 | *CSTF2T* |
|  | cg02366575 | 2502 | 10.25 | 2.27 | 6.17E-06 | 9.3 | 1 | 91966307 | *CDC7* |
|  | cg12813441 | 2506 | -1.61 | 0.36 | 7.88E-06 | 0 | 2 | 55239331 | *RTN4* |
|  | cg21950196 | 2512 | 2.45 | 0.55 | 8.51E-06 | 0 | 1 | 228327428 | *GUK1* |

^a^ Beta coefficient from the regression indicating the change in IQ score standard deviations per 100% methylation change. Models were adjusted also for age at testing, sex, maternal age at delivery, maternal education, birthweight, gestational age, maternal smoking status during pregnancy, parity, batch covariates and cell proportions

^b^ Unadjusted p-value

^c^ Heterogeneity statistics

**Supplementary Table ST7.** Association estimates at top sites from the main model EWAS meta-analyses across all sensitivity models.

|  |  | **Main model** | | | **Paternal education** | | | **Maternal IQ** | | | **Genetic PCs** | | |
| --- | --- | --- | --- | --- | --- | --- | --- | --- | --- | --- | --- | --- | --- |
|  | **CpG site** | **Beta** | **S.E.** | **P-value** | **Beta** | **S.E.** | **P-value** | **Beta** | **S.E.** | **P-value** | **Beta** | **S.E.** | **P-value** |
| **Overall** | cg05827775 | -3.05 | 0.64 | 1.84E-06 | -2.95 | 0.66 | 7.73E-06 | -1.42 | 1.11 | 0.20 | -2.93 | 0.84 | 0.0005 |
|  | cg00213080 | -2.44 | 0.51 | 1.89E-06 | -2.48 | 0.52 | 1.48E-06 | -1.39 | 1.25 | 0.27 | -2.44 | 0.62 | 7.53E-05 |
|  | cg26599274 | 10.83 | 2.30 | 2.46E-06 | 10.58 | 2.36 | 7.17E-06 | 4.43 | 4.27 | 0.30 | 11.97 | 2.92 | 4.09E-05 |
|  | cg23789148 | 2.03 | 0.44 | 3.88E-06 | 2.12 | 0.46 | 4.60E-06 | 2.07 | 1.05 | 0.05 | 2.34 | 0.51 | 4.59E-06 |
|  | cg18622281 | 3.96 | 0.86 | 3.91E-06 | 3.90 | 0.87 | 7.76E-06 | 2.38 | 1.57 | 0.13 | 3.50 | 1.13 | 0.002 |
|  | cg00573504 | 3.71 | 0.81 | 4.98E-06 | 3.52 | 0.83 | 2.11E-05 | 2.96 | 2.48 | 0.23 | 2.93 | 0.94 | 0.002 |
|  | cg09535605 | 5.57 | 1.22 | 5.09E-06 | 5.13 | 1.23 | 2.94E-05 | 3.16 | 2.36 | 0.18 | 5.77 | 2.09 | 0.006 |
|  | cg21735491 | -9.38 | 2.09 | 6.77E-06 | -9.20 | 2.08 | 9.38E-06 | -9.13 | 7.10 | 0.20 | -10.75 | 2.85 | 0.0002 |
|  | cg18075761 | -23.68 | 5.36 | 9.84E-06 | -21.74 | 5.43 | 6.30E-05 | -11.79 | 12.80 | 0.36 | -20.66 | 6.98 | 0.003 |
| **Verbal** | cg03568675 | -3.00 | 0.59 | 4.36E-07 | -2.8 | 0.64 | 9.28E-06 | -0.81 | 1.31 | 0.54 | -2.65 | 0.70 | 0.0002 |
|  | cg12361663 | 36.38 | 7.70 | 2.30E-06 | 35.71 | 7.67 | 3.22E-06 | 35.54 | 19.19 | 0.06 | 37.28 | 11.47 | 0.001 |
|  | cg17223866 | 5.21 | 1.12 | 3.14E-06 | 3.92 | 1.19 | 0.001 | 7.38 | 2.95 | 0.01 | 5.30 | 1.27 | 3.05E-05 |
|  | cg11005998 | 2.94 | 0.64 | 3.87E-06 | 2.29 | 0.67 | 0.0006 | 2.70 | 2.43 | 0.27 | 2.41 | 0.71 | 0.0007 |
|  | cg10620273 | 22.72 | 5.00 | 5.65E-06 | 22.04 | 4.97 | 9.19E-06 | 5.49 | 10.12 | 0.59 | 10.62 | 8.13 | 0.19 |
|  | cg16047144 | -13.16 | 2.92 | 6.33E-06 | -12.01 | 2.99 | 5.88E-05 | -11.78 | 6.20 | 0.06 | -13.13 | 4.01 | 0.001 |
| **Non-verbal** | cg04783204 | 8.90 | 1.91 | 3.11E-06 | 9.76 | 1.99 | 8.97E-07 | 6.76 | 2.32 | 0.004 | 9.34 | 2.22 | 2.63E-05 |
|  | cg04229103 | 6.34 | 1.36 | 3.31E-06 | 6.58 | 1.42 | 3.74E-06 | 4.99 | 1.63 | 0.002 | 5.60 | 1.58 | 0.0004 |
|  | cg00573504 | 3.38 | 0.73 | 3.32E-06 | 3.39 | 0.76 | 7.53E-06 | 1.47 | 1.17 | 0.21 | 2.57 | 0.80 | 0.001 |
|  | cg25990848 | 2.07 | 0.45 | 3.59E-06 | 2.18 | 0.46 | 2.45E-06 | 2.92 | 1.14 | 0.01 | 2.04 | 0.49 | 3.59E-05 |
|  | cg08529049 | -4.95 | 1.08 | 4.30E-06 | -4.25 | 1.17 | 0.0003 | -5.39 | 1.45 | 0.0002 | -4.06 | 1.19 | 0.0007 |
|  | cg03332597 | 3.83 | 0.84 | 4.80E-06 | 4.30 | 0.86 | 6.49E-07 | 2.55 | 1.04 | 0.02 | 3.70 | 0.96 | 0.0001 |

**Supplementary Table ST8.** Replication of sites found in Marioni et al. (2018): results for the top two sites across main models and results for the replicated at Bonferroni-adjusted p-value<0.05

|  | **CpG site** | **Beta**^a^ | **S.E.** | **P-value** ^b^ | **I^2^** ^c^ | **Direction of effect in Marioni et al.** |
| --- | --- | --- | --- | --- | --- | --- |
| **Overall** | cg12507869 | 0.55 | 1.22 | 0.65 | 29.0 | - |
|  | cg21450381 | 1.37 | 1.02 | 0.18 | 0 | - |
| **Verbal** | cg12507869 | 1.04 | 1.24 | 0.40 | 46.6 | - |
|  | cg21450381 | 1.45 | 1.03 | 0.16 | 0 | - |
| **Non-verbal** | cg12507869 | -0.87 | 0.92 | 0.34 | 34.5 | - |
|  | cg21450381 | NA | NA | NA | NA | - |
| **Non-verbal**  **(replicated site)** | cg17759224 | 1.17 | 0.34 | 0.00065 | 0 | + |

^a^ Beta coefficient from the regression indicating the change in IQ score standard deviations per 100% methylation change. Models were adjusted also for age at testing, sex, maternal age at delivery, maternal education, birthweight, gestational age, maternal smoking status during pregnancy, parity, batch covariates and cell proportions

^b^ Unadjusted p-value

^c^ Heterogeneity statistics

NA= not available in the meta-analysis

**Supplementary Table ST9.** Replication of sites found in Karlsson-Linner et al. (2018): results for top two sites across main models and results for the replicated at Bonferroni-adjusted p-value<0.05

|  | **CpG site** | **Beta**^a^ | **S.E.** | **P-value** ^b^ | **I^2^** ^c^ | **Direction of effect in Karlsson-Linner et al. 2017** |
| --- | --- | --- | --- | --- | --- | --- |
| **Overall** | cg01940273 | -1.06 | 0.61 | 0.08 | 0 | + |
|  | cg03636183 | 0.62 | 0.44 | 0.15 | 0 | + |
|  | cg05575921 | -0.75 | 0.47 | 0.11 | 0 | + |
|  | cg05951221 | -0.07 | 0.88 | 0.94 | 0 | + |
|  | cg06126421 | NA | NA | NA | NA | + |
|  | cg12803068 | 0.02 | 0.21 | 0.94 | 47.7 | - |
|  | cg21161138 | 0.19 | 0.32 | 0.56 | 0 | + |
|  | cg21566642 | 0.65 | 0.80 | 0.42 | 0 | + |
|  | cg22132788 | 0.03 | 0.44 | 0.94 | 45.2 | - |
| **Verbal** | cg01940273 | -0.59 | 0.61 | 0.33 | 0 | + |
|  | cg03636183 | 0.52 | 0.44 | 0.24 | 0 | + |
|  | cg05575921 | -0.64 | 0.48 | 0.18 | 0 | + |
|  | cg05951221 | 0.23 | 0.89 | 0.79 | 0 | + |
|  | cg06126421 | NA | NA | NA | NA | + |
|  | cg12803068 | 0.22 | 0.21 | 0.29 | 33.8 | - |
|  | cg21161138 | -0.04 | 0.33 | 0.90 | 0 | + |
|  | cg21566642 | 0.25 | 0.81 | 0.76 | 0 | + |
|  | cg22132788 | 0.67 | 0.44 | 0.13 | 0 | - |
| **Non-verbal** | cg01940273 | -0.38 | 0.55 | 0.48 | 14.0 | + |
|  | cg03636183 | 0.70 | 0.41 | 0.09 | 0 | + |
|  | cg05575921 | -0.21 | 0.40 | 0.61 | 29.5 | + |
|  | cg05951221 | 0.74 | 0.67 | 0.27 | 2.8 | + |
|  | cg06126421 | NA | NA | NA | NA | + |
|  | cg12803068 | -0.29 | 0.19 | 0.13 | 0 | - |
|  | cg21161138 | 0.14 | 0.32 | 0.65 | 0 | + |
|  | cg21566642 | 0.85 | 0.65 | 0.19 | 0 | + |
|  | cg22132788 | NA | NA | NA | NA |  |

^a^ Beta coefficient from the regression indicating the change in IQ score standard deviations per 100% methylation change. Models were adjusted also for age at testing, sex, maternal age at delivery, maternal education, birthweight, gestational age, maternal smoking status during pregnancy, parity, batch covariates and cell proportions

^b^ Unadjusted p-value

^c^ Heterogeneity statistics

NA= not available in the meta-analysis

**Supplementary Table S10**. Differentially methylated region from the meta-analysis at Bonferroni-corrected p<0.05

| **CpG site** | **N** | **Beta**^a^ | **S.E.** | **P-value** ^b^ | **I^2^** ^c^ | **Chr.** | **Position** | **Gene** ^d^ |
| --- | --- | --- | --- | --- | --- | --- | --- | --- |
| cg03395511 | 1604 | 0.26 | 0.17 | 0.121 | 0 | chr6 | 291903 | *DUSP22* |
| cg07332563 | 1862 | 0.34 | 0.13 | 0.009 | 0 | chr6 | 291687 | *DUSP22* |
| cg15383120 | 1804 | 0.27 | 0.12 | 0.029 | 0 | chr6 | 291909 | *DUSP22* |
| cg18110333 | 1862 | 0.26 | 0.10 | 0.012 | 5.3 | chr6 | 292329 | *DUSP22* |
| cg21548813 | 1804 | 0.26 | 0.13 | 0.041 | 0 | chr6 | 291882 | *DUSP22* |

^a^ Beta coefficient from the regression indicating the change in IQ score standard deviations per 100% methylation change. Models were adjusted also for age at testing, sex, maternal age at delivery, maternal education, birthweight, gestational age, maternal smoking status during pregnancy, parity, batch covariates, cell proportions and maternal IQ

^b^ Unadjusted p-value

^c^ Heterogeneity statistics

|  | **Prefrontal cortex** | | **Entorhinal cortex** | | **Cerebellum** | | **Superior temporal gyrus** | |
| --- | --- | --- | --- | --- | --- | --- | --- | --- |
| **CpG** | **r** | **p-value** | **r** | **p-value** | **r** | **p-value** | **r** | **p-value** |
| cg03395511 | 0.981 | 3.83e-53 | 0.965 | 5.05e-42 | 0.984 | 6.30e-57 | 0.975 | 5.60e-47 |
| cg07332563 | 0.946 | 5.55e-37 | 0.940 | 7.93e-34 | 0.951 | 4.76e-39 | 0.927 | 3.38e-31 |
| cg15383120 | 0.971 | 1.67e-46 | 0.939 | 9.17e-34 | 0.961 | 2.26e-42 | 0.937 | 2.59e-33 |
| cg18110333 | 0.986 | 6.57e-58 | 0.981 | 7.78e-51 | 0.986 | 4.74e-58 | 0.970 | 4.82e-44 |
| cg21548813 | 0.986 | 5.33e-58 | 0.975 | 7.80e-47 | 0.988 | 9.75e-61 | 0.986 | 2.11e-55 |

**Supplementary Table S11**. Correlation of DNA methylation levels between brain and blood in adult samples (N=71-75) in the *DSUP22* differentially methylated region (<https://epigenetics.essex.ac.uk/bloodbrain/>).

**Supplementary Table S12.** Correlation between DNA methylation and gene expression at the DUSP22 differentially-methylated region (https://genenetwork.nl/biosqtlbrowser/).

| **CpG** | **P-value** | **CpG**  **Chr.** | **CpG Chr.**  **Position** | **Probe** | **Probe**  **Chr.** | **Probe Chr.**  **position** | **Z-score** |
| --- | --- | --- | --- | --- | --- | --- | --- |
| cg03395511 | 2.63E-154 | 6 | 291855 | ENSG00000112679 | 6 | 292097 | 26.46 |
| cg07332563 | 4.11E-163 | 6 | 291639 | ENSG00000112679 | 6 | 292097 | 27.22 |
| cg15383120 | 4.4E-160 | 6 | 291861 | ENSG00000112679 | 6 | 292097 | 26.96 |
| cg18110333 | 3.83E-155 | 6 | 292329 | ENSG00000112679 | 6 | 292097 | 26.53 |
| cg21548813 | 2.44E-157 | 6 | 291834 | ENSG00000112679 | 6 | 292097 | 26.72 |
