## supplementary material for "Meta-analysis of epigenome-wide associations between DNA methylation at birth and childhood cognitive skills"

**Supplementary Material and Methods**

ALSPAC

The Avon Longitudinal Study of Parents and Children (ALSPAC) ^1, 2^ is a prospective pregnancy cohort based in the United Kingdom. Pregnant women resident in Avon, UK with expected dates of delivery between 1^st^ April 1991 and 31^st^ December 1992 were invited to take part in the study. An initial number of 14,541 women returned questionnaire or attended a clinic subsequently. Of these pregnancies there were 13,988 children who were alive at 1 year of age. The study website contains details of all the available through a fully searchable data dictionary and variable search tool (<http://www.bristol.ac.uk/alspac/researchers/our-data/>). Ethical approval for the study was obtained from the ALSPAC Ethics and Law Committee and the Local Research Ethics Committees. Consent for biological samples has been collected in accordance with the Human Tissue Act (2004). Informed consent for the use of data collected via questionnaires and clinics was obtained from participants following the recommendations of the ALSPAC Ethics and Law Committee at the time.

DNA methylation was measured in cord blood from 1018 of these children as part of the Accessible Resource for Integrative Epigenomic Studies (ARIES) ^3^. Briefly, following DNA extraction samples were bisulphite converted using the Zymo EZ DNA Methylation™ kit (Zymo, Irvine, CA, USA). Following conversion, genome-wide methylation was measured using the Illumina Infinium HumanMethylation450 (HM450) BeadChip. The arrays were scanned using an Illumina IScan, with initial quality review using GenomeStudio. Pre-processing and normalization were done using the meffil R package. Full details of the pre-processing and normalisation has been described previously ^4^. Surrogate variable analysis using the *sva* R package ^5^ identified 10 surrogate variables that were then entered in all models to correct for batch.

Intelligent quotient (IQ) was assessed in the children participants when they were approximately 8 years of age using a short form of the Wechsler Intelligence Scale for Children WISC-III UK ^6^ which was administered by a trained psychologist. The test consisted of ten subtests, which were then combined to obtain a raw verbal score (information, similarities, arithmetic, vocabulary and comprehension subtests) and a raw performance score (picture completion, coding, picture arrangement, block design, and object assembly). Age-scaled scores for full-scale, performance and verbal IQ were then computed using the look-up tables in the WISC manual. Prorating was then performed according to the WISC manual instructions. Information on maternal and paternal education was derived by questionnaires administered to the pregnant women at recruitment and was categorised as having at least a university degree compared to less education than university degree.

CHAMACOS

The Center for the Health Assessment of Mothers and Children of Salinas (CHAMACOS) study is a longitudinal birth cohort study of the effects of exposure to pesticides and environmental chemicals on the health and development of Mexican-American children living in the agricultural region of Salinas Valley, CA (USA). Detailed description of the CHAMACOS cohort has previously been published ^7, 8^.Briefly, 601 pregnant women were enrolled in 1999- 2000 at community clinics and 527 liveborn singletons were born. Follow up visits occurred at regular intervals throughout childhood.

DNA methylation was measured in DNA isolated from the cord blood of 367 newborns. The DNA samples were bisulfite converted using Zymo Bisulfite Conversion Kits (Zymo Research, Irvine, CA), whole genome amplified, enzymatically fragmented, purified, and applied to the 450K BeadChips (Illumina, San Diego, CA) according to manufacturer protocol. 450K BeadChips were handled by robotics and analysed using the Illumina Hi_Scan system. Probe signal intensities were extracted by Illumina GenomeStudio software (version XXV2011.1, Methylation Module 1.9) methylation module and background subtracted. QA/QC was performed systematically by assessment of assay repeatability batch effects using 38 technical replicates, and data quality established as previously described ^9^. Quality was also ensured by only retaining samples where 95 % of sites assayed had detection p> 0.01. The same threshold (95% detection at p>0.01) was imposed to CpGs as well (n= 460 removed). Sites with annotated probe SNPs and with common SNPs (minor allele frequency >5%) within 50bp of the target identified in the MXL (Mexican ancestry in Los Angeles, California) HapMap population were excluded from analysis (n=49,748). This left a total of 435,369 CpGs in the analysis. Color channel bias, batch effects and difference in Infinium chemistry were minimized by application of ASMN algorithm ^9^, followed by BMIQ normalization ^10^.

Childhood intelligence score was assessed at approximately 7 years of age using the Wechsler Intelligence Scale for Children , 4^th^ edition (WISC-IV) ^11^. The scores used in the analysis were the full-scale score, verbal comprehension index and perceptual reasoning index.

Maternal and paternal education covariates were treated as categorical, with two levels: less than a completed high school education, having completed high school education, or beyond. Batch effects were accounted for by adjusting for 450K plate as additional covariate. Maternal IQ was assessed when the children were 6 months using the Peabody Picture Vocabulary Test ^12^.

Generation R Study

Data for the current study were drawn from a European subsample of the Generation R Study. The Generation R Study is a prospective population-based cohort. Pregnant women with an expected delivery date between April 2002 and January 2006 residing in the municipality of Rotterdam, the Netherlands, were invited to enroll in the study. In total, 9778 pregnant women had 9749 live-born children. DNA methylation was assessed in1396 children at birth. A more detailed description of the Generation R Study can be found elsewhere ^13, 14^. The Generation R Study is conducted in accordance with the World Medical Association Declaration of Helsinki and has been approved by the Medical Ethics Committee of the Erasmus Medical Center, Rotterdam. Written informed consent was obtained for all participants.

DNA was extracted using the salting-out method from cord blood. 500 ng of DNA per sample underwent bisulfite conversion using the EZ-96 DNA methylation kit (Shallow) (Zymo Research Corporation, Irvine, USA). Samples were plated onto 96-well plates in no specific order. The bisulfite-converted DNA was then processed with the Illumina Infinium HumanMethylation450 BeadChip (Illumina Inc., San Diego, USA). Pre-processing and normalisation was performed according to the CPACOR ^15^ workflow using R software. In detail, the idat files were read using the minfi R package ^16^. Probes that had a detection p-value above background ≥ 1E-16 were set to missing per array. Next, the intensity values were stratified by autosomal and non-autosomal probes and quantile normalised for each of the six probe type categories separately. Beta values were calculated as proportion of methylated intensity value on the sum of methylated+unmethylated+100 intensities. Arrays with failed bisulphite conversion, hybridisation, extension or mismatch between reported sex of the proband and sex determined by the chr X and Y probe intensities were excluded. Only arrays with a call rate >95% per samples were retained for further analysis, leading to a final sample of 1396 participants. Batch effects were corrected by adjusting for sample plate (n=28, dummy coding).

Cognitive ability was measured at the Generation R research center when the children were six years old. The Generation R Study features a high proportion of children with non-Dutch national origin. To avoid potential biases as consequence of non-native language backgrounds, only non-verbal tests using the Snijders‐Oomen Niet‐verbale intelligentie test 2.5-7 ^17^ were administrated. We tested spatial and abstract reasoning with the “Mosaics” and “Categories” subtests, respectively. We then transformed the obtained raw scores into nonverbal IQ scores. Maternal education at delivery was categorized into primary, secondary phase one, secondary phase two, higher phase one and higher phase two, and analyzed as continuous variable. Gestational age was estimated with an ultrasound examination. We measured maternal IQ when the children were six years old at the research center using the first set of the Raven’s Advanced Progressive Matrices Test ^18^. Genetic ancestry was obtained by a multi-dimensional scaling analysis using genome-wide SNP data obtained with the Illumina HumanHap 610 or 660 Quad chips, see Medina-Gomez et al ^19^.

EDEN

The EDEN (Etude des Déterminants pré et post natals du développement et de la santé de l′Enfant) study is a prospective birth cohort study (https://eden.vjf.inserm.fr/), which has been described in detail elsewhere ^20^. Pregnant women seen for a prenatal visit at the departments of Obstetrics and Gynecology of the University Hospital of Nancy and Poitiers before their twenty-fourth wek of amenorrhea were invited to participate. Enrolment started in February 2003 in Poitiers and September 2003 in Nancy; it lasted 27 months in each centre. Among eligible women, 55% (n=2002) accepted to participate.

Immediately after delivery, cord blood samples were collected by research midwives from 1367 consenting cohort participants. To prevent any contamination with maternal blood, the cord was doubly clamped immediately after birth (vaginal delivery) or after extraction of the fetus through the uterine incision (elective cesarean section); repeatedly rinsed and venous cord blood serum was sampled between the 2 clamps. Cord blood samples were centrifuged within 24 hours of collection. The serum was separated and samples were stored at −80°C.Cord blood samples were collected from all consenting cohort participants and DNA was extracted using the QIAamp blood kit (Qiagen or equivalent protocols), followed by precipitation-based concentration using GlycoBlue (Ambion). DNA concentration was determined by Nanodrop measurement and Picogreen quantification. 500 ng of DNA was bisulphite-converted using the EZ 96-DNA methylation kit (Zymo Research), following the manufacturer’s standard protocol. After verification of the bisulphite conversion step using Sanger Sequencing, genome-wide DNA methylation was measured using the Illumina Infinium HumanMethylation450 BeadChip. After normalization of the concentration, the samples were randomized to avoid batch effects, and all paired samples were hybridized on the same chip. Standard male and female DNA samples were included in this step as control samples. DNA methylation data were pre-processed in R with the Bioconductor package minfi ^16^, using the original IDAT files extracted from the HiScanSQ scanner. Samples that did not provide significant methylation signals in more than 10% of probes (detection P = 0.01) were excluded from further analysis. Samples were also excluded in cases of low staining efficiency, low single base extension efficiency, low stripping efficiency of DNA from probes after single base extension, poor hybridization performance, poor bisulfite conversion and high negative control probe staining. Further, we used the 65 SNP probes to check for concordances between paired DNA samples from the sample individual and assessed the methylation distribution of the X-chromosome to verify gender. Paired samples with Pearson correlation coefficients <0.9 were regarded as sample mix-ups and were excluded from the study. In probe filtering, we excluded probes on sex chromosomes, probes that mapped on multi-loci, the 65 random SNPs assay and probes that contained SNPs at the target CpG sites with a minor allele frequency >10% ^21^. The allele frequencies of a list of SNPs were obtained from 1000 Genomes, release 20110521 for CEU population. Finally, we implemented “DASEN” to perform signal correction and normalization ^22^. After quality control, 165 samples and 439,306 autosomal probes remained. From these, we selected 158 samples from the population of randomly selected children for further analysis based on completeness of data for study covariates.

The Wechsler Preschool and Primary Scale of Intelligence (WPPSI) 3rd Edition was administered by trained psychologists when the children were aged 5 years. The core subsets of the battery were assessed to obtain the composite scores of verbal IQ, performance IQ, and full-scale IQ, which were age-normed in accordance with standard procedures ^23^. Maternal education was based on the self-reported highest school diploma obtained and categorised in 3 classes: 0 if obtained primary education or pre-professional courses (age 16 or younger); 1 if obtained a high school degree (baccalauréat, age 17-18); 2 if obtained a tertiary education degree (bachelor degree or above).

INMA

The INMA -- INfancia y Medio Ambiente -- (Environment and Childhood) Project is a network of birth cohorts in Spain aimed to studying the role of environmental pollutants in air, water and diet during pregnancy and early childhood in relation to child growth and development

(<http://www.proyectoinma.org/>) ^24^. Data for this study comes from INMA Sabadell sub-cohort.

A full roster of the INMA Project Investigators

can be found at <http://www.proyectoinma.org/presentacion-inma/listado-investigadores/en_listado-investigadores.html>.

Cord blood DNA was extracted using the Chemagen kit (Perkin Elmer) and its concentration was determined by NanoDrop spectrophotometer (Thermo Scientific) and with the Quant-iT PicoGreen dsDNA Assay Kit (Life Technologies). Methylation data was produced in two

different laboratories as part of two different projects: in the Genome Analysis Facility of the University Medical Center Groningen in the Netherlands, and in the Bellvitge Biomedical Research Institute (IDIBELL, Barcelona), both using the recommended Illumina

protocol for the Infinium HumanMethylation450 beadchip. Briefly, 500 ng of DNA was bisulphite-converted using the EZ 96-DNA methylation kit following the manufacturer’s standard protocol, and DNA methylation measured using the Illumina Infinium HumanMethylation450 beadchip. DNA methylation data were pre-processed using the minfi R package ^15^. During quality control two samples with bad overall quality or with low detection p-value according to the output of the MethylAid R package ^25^ and three samples whose sex was wrongly predicted using the shinyMethyl R package ^26^ were removed. Eighteen samples with a call rate lower than 98% detection (p-value threshold to 10^-16^) were also removed as previously recommended ^15^. Functional normalisation was applied. The correlation between SNP probes in replicates samples was checked and samples with discordance were discarded. 7,136 probes with a call rate lower than 95% were also removed. ComBat was applied to remove batch effect ^27^. Finally, duplicated samples were removed. The final dataset consisted of 319 samples with cognition scores available.

Childhood cognition was assessed using the McCarthy Scales of Children’s Abilities ^28^ at approximately 5 years of age. The general cognitive index, the verbal index and the perceptual-performance index were used as measures of overall, verbal and non-verbal cognitive skills, respectively.

Maternal and paternal education was grouped in 3 categories: low (primary or less), medium (secondary), high (university). Maternal IQ was estimated using the similarities subtest from the Weschler Adult Intelligence-Third Edition (WAIS-III) ^29^.

POSEIDON

In total 410 pregnant women were recruited about 4–8 weeks prior to delivery from hospitals in the Rhine-Neckar Region of Germany, to take part in a longitudinal study on perceived stress and child development and health (Pre-, Peri-, and Postnatal Stress: Epigenetic impact on Depression; POSEIDON) ^30^.

Mothers were included if they were 16–45 years old, German-speaking and presumably the child’s main caregiver. Maternal Exclusion criteria were a diagnosis of hepatitis B, hepatitis C or human immunodeficiency virus, current psychiatric disorder requiring inpatient treatment, history or current diagnosis of schizophrenia or psychotic disorder, substance dependency other than nicotine during pregnancy. Exclusion criteria for newborn children were birth before 30 weeks of pregnancy, birthweight less than 1,500 g, multiple birth, congenital disease, malformation, deformation or chromosomal abnormality.  Data were collected during third trimester of pregnancy (T1), a few days after childbirth (T2), six months postpartum (T3) and 45 months postpartum (T4). The last assessment (T4) took place between August 2014 and January 2017(4).
Intensity data were extracted from raw data (idat) files using an updated version of the pipeline published by Lehne et al ^15^. Intensity data were then quantile normalized within subsets of probe types prior to converting to beta values. Samples were excluded in case of insufficient DNA quality, insufficient bisulfite conversion, failure in detection (detection P-value > 0.01 at more than 1% of positions), or sex-mismatch between phenotype and methylation data. QC steps involved exclusion of probes with detection p-value threshold (positions/sites) > 0.01 or with a call rate < 95%, exclusion of sex chromosomes. The first 10 principle components of the control probes were included as covariates to account for batch effects.

Samples were genotyped on the Illumina PsychChip array (Illumina, San Diego, CA) using the PsychChip 15048346 B manifest. Population structure was determined based on a SNP set filtered for high quality (HWE p > .02, MAF > .20, missingness = 0), and LD pruning (r² = .1). Principal components were generated to control for population stratification and were included in the EWAS for models 10-12.

Cognitive skills were measured using the Wechsler Preschool and Primary Scale of Intelligence (WPPSI) at approximately 4 years of age. Full-scale, verbal and performance IQ scores were used as overall, verbal and non-verbal cognitive skills.

Parents’ education, maternal age and smoking were measured by questionnaires administered during the third pregnancy trimester. Maternal and paternal education were measured in years of schooling. Briefly, during the third pregnancy trimester mothers were asked for their highest educational level, highest training qualification and highest degree (college, academy, or university). The ISCED 1997 levels were used to transform the highest education attainment in our sample. Afterwards the ISCED levels were transformed into US years of schooling.

PREDO

The Prediction and Prevention of Preeclampsia and Intrauterine Growth (PREDO) is a prospective, multi-centre study based in Finland that recruited 5332 women with a single intrauterine pregnancy between 2005 and 2009 and their children ^31^. Of these, 4777 were eligible, consented to participate and resulted to live births. A subsample of 285 children participants had cord blood DNA methylation, cognitive skills data and main covariates available. DNA methylation was measured using the Illumina 450K Methylation microarrays. All samples were randomized on 96-well plates based on gender and maternal risk factors. The quality control pipeline was set up using the R-package *minfi* (<https://www.r-project.org>). Median intensities outliers, samples with discordance between phenotypic sex and estimated sex, and samples contamination with maternal DNA Methylation wre excluded. Beta-values were normalized using the funnorm function. Furthermore, any CpGs with a detection p-value > 0.01 in at least 25% of the samples were excluded. After normalization, two batches, i.e., slide and well, were significantly associated and were removed iteratively using the Combat function in the *sva* R package ^5^.

Cognitive skills were measured using the WISC-IV test ^11^ at approximately 8 years of age. Infant sex, maternal age at delivery, maternal and paternal education, smoking during pregnancy, and parity data were derived from the Finnish Medical Birth Register. The first two principal components derived from genome-wide level genotypes were included in the PC-adjusted model.

Project Viva

Project Viva is a prospective pre-birth cohort study of mother–child pairs recruited between 1999 and 2002 during the mothers’ first prenatal visits at Atrius Harvard Vanguard Medical Associates, a multi-specialty medical group practice in Massachusetts, United States ^32^. Exclusion criteria included multiple gestation, inability to answer questions in English, gestational age ≥22 weeks at recruitment and plans to move away before delivery. Mothers provided written informed consent at recruitment and at postpartum follow-up visits. The Institutional Review Board of Harvard Pilgrim Health Care reviewed and approved all study protocols.

Cord blood DNA was extracted and bisulfite treated. DNA methylation was measured using the Illumina Infinium HumanMethylation450 BeadChip (Illumina Inc., San Diego, USA). Data pre-processing involved the removal of failed samples (as indicated in the Illumina sample-sheet crossed referenced with *minfi*), removal or replicates and elimination of non-CpG probes. The data were then checked for gender mismatch using the X and Y chromosomes. CpGs with low P-values were identified and flagged. Samples with mismatch based on genotype were removed. Raw methylation values were Noob-adjusted ^33^. Further adjustment for BMIQ based on probe type (I vs II) was performed. Finally, batch correction was performed using ComBat to adjust for sample plate.

Childhood cognition was assessed during in-person mid-childhood follow-up visits by research assistants either in the child’s home or at the central research office at approximately 8 years of age. Total IQ was assessed by administering the Wide Range Assessment of Visual Motor Ability (WRAVMA) test ^34^, which evaluates three domains of visual motor development: visual-spatial (matching test), visual-motor (drawing test), and fine motor skills (pegboard test). The three tests are used to generate a total standard score. This test has norms for children aged 3 years and older and it is moderately correlated with IQ (r~0.60). Assessments were double-scored using published scoring guidelines and supplementary guidelines developed by a paediatric neuropsychologist to ensure consistency among scorers. Verbal and non-verbal intelligence were measured using the Kaufman Brief Intelligence Test 2^nd^ edition (KBIT-II) ^35^. The verbal subtest contains two items types (verbal knowledge and riddles) that measure crystallised intelligence; the non-verbal subtest includes matrices that measures fluid reasoning. The two subtests provide an IQ composite that correlates with full-length measures of intelligence such as the WISC-III (r=0.63). Maternal intelligence was assessed using the KBIT-II during the same visit.

**ACKNOWLEDGEMENTS**

We are extremely grateful to all the families who took part in the ALSPAC study, the midwives for their help in recruiting them, and the whole ALSPAC team, which includes interviewers, computer and laboratory technicians, clerical workers, research scientists, volunteers, managers, receptionists and nurses. We thank all parents and children for taking part in the POSEIDON study. We thank the Erasmus Medical Center, the Erasmus University Rotterdam, Faculty of Social Sciences, the Municipal Health Service Rotterdam area, the Rotterdam Homecare Foundation, Rotterdam and the Stichting Trombosedienst & Artsenlaboratorium Rijnmond (STAR-MDC), Rotterdam, for conducting the Generation R study. We gratefully acknowledge the contribution of general practitioners, hospitals, midwives and pharmacies in Rotterdam. The generation and management of the Illumina 450K methylation array data (EWAS data) for the Generation R Study was executed by the Human Genotyping Facility of the Genetic Laboratory of the Department of Internal Medicine, Erasmus MC, the Netherlands. We thank Mr. Michael Verbiest, Ms. Mila Jhamai, Ms. Sarah Higgins, Mr. Marijn Verkerk and Dr. Lisette Stolk for their help in creating the EWAS database. We thank Dr. A.Teumer for his work on the quality control and normalization scripts. INMA researchers would like to thank all the participants for their generous collaboration. INMA researchers are grateful to Silvia Fochs, Nuria Pey, and Muriel Ferrer for their assistance in contacting the families and administering the questionnaires. A full roster of the INMA Project Investigators can be found at <http://www.proyectoinma.org/presentacion-inma/listadoinvestigadores/en_listado-investigadores.html>. We are grateful to the CHAMACOS participants and field staff for their contributions. We are indebted to the EDEN participating families and the midwife research assistants (L. Douhaud, S. Bedel, B. Lortholary, S. Gabriel, M. Rogeon, M. Malinbaum) for data collection. We thank the EDEN mother-child cohort study group (I. Annesi-Maesano, J.Y Bernard, J. Botton, M.A. Charles, P. Dargent-Molina, B. de Lauzon-Guillain, P. Ducimetière, M. de Agostini, B. Foliguet, A. Forhan, X. Fritel, A. Germa, V. Goua, R. Hankard, B. Heude, M. Kaminski, B. Larroque†, N. Lelong, J. Lepeule, G. Magnin, L. Marchand, C. Nabet, F. Pierre, R. Slama, M.J. Saurel-Cubizolles, M. Schweitzer, O. Thiebaugeorges). The PREDO study would not have been possible without the dedicated contribution of the PREDO study group members: E Hamäläinen, E Kajantie, H Laivuori, PM Villa, A-K Pesonen, A Aitokallio-Tallberg, A-M Henry, VK Hiilesmaa, T Karipohja, R Meri, S Sainio, T Saisto, S Suomalainen-Konig, V-M Ulander, T Vaitilo (Department of Obstetrics and Gynaecology, University of Helsinki and Helsinki University Central Hospital, Helsinki, Finland), L Keski-Nisula, Maija-Riitta Orden (Kuopio University Hospital, Kuopio Finland), E Koistinen, T Walle, R Solja (Northern Karelia Central Hospital, Joensuu, Finland), M Kurkinen (Päijät-Häme Central Hospital, Lahti, Finland), P.Taipale. P Staven (Iisalmi Hospital, Iisalmi, Finland), J Uotila (Tampere University Hospital, Tampere, Finland). We thank all the PREDO children and their parents for their enthusiastic participation. We also thank all the research nurses, research assistants, and laboratory personnel involved in the PREDO study.

**ETHICAL STATEMENT**

Ethical approval for the ALSPAC study was obtained from the ALSPAC Ethics and Law Committee and the Local Research Ethics Committees. Consent for biological samples has been collected in accordance with the Human Tissue Act (2004). Informed consent for the use of data collected via questionnaires and clinics was obtained from participants following the recommendations of the ALSPAC Ethics and Law Committee at the time. The POSEIDON study protocol was approved by the Ethics Committee of the Medical Faculty Mannheim of the University of Heidelberg. The PREDO study was conducted in accordance with the Declaration of Helsinki. The Generation R study has been approved by the Medical Ethical Committee of the Erasmus MC, University Medical Center Rotterdam (MEC 198.782/2001/31). Written informed consent was obtained from all adult participants. The INMA study has been approved by the Ethical Committee of the Institut Municipal d’Investigació Mèdica, Barcelona

and written consent was obtained from participating parents. Project Viva mothers provided written informed consent and protocols were approved by Institutional Review Board at Harvard Pilgrim Health Care. CHAMACOS study protocols were approved by the University of California, Berkeley Committee for Protection of Human Subjects and written informed consent was obtained from all mothers; oral assent was obtained from children beginning at age 7, and written assent at age 12. The EDEN study has been approved by the ethical committees « Comité Consultatif pour la Protection des Personnes dans la Recherche Biomédicale », Le Kremlin-Bicêtre University hospital, and «Commission Nationale de l’Informatique et des Libertés». The Ethics Committees of the Helsinki and Uusimaa Hospital District and the participating hospitals approved the study protocol. All participating women signed written informed consents.
