## supplementary figures for "Meta-analysis of epigenome-wide associations between DNA methylation at birth and childhood cognitive skills"

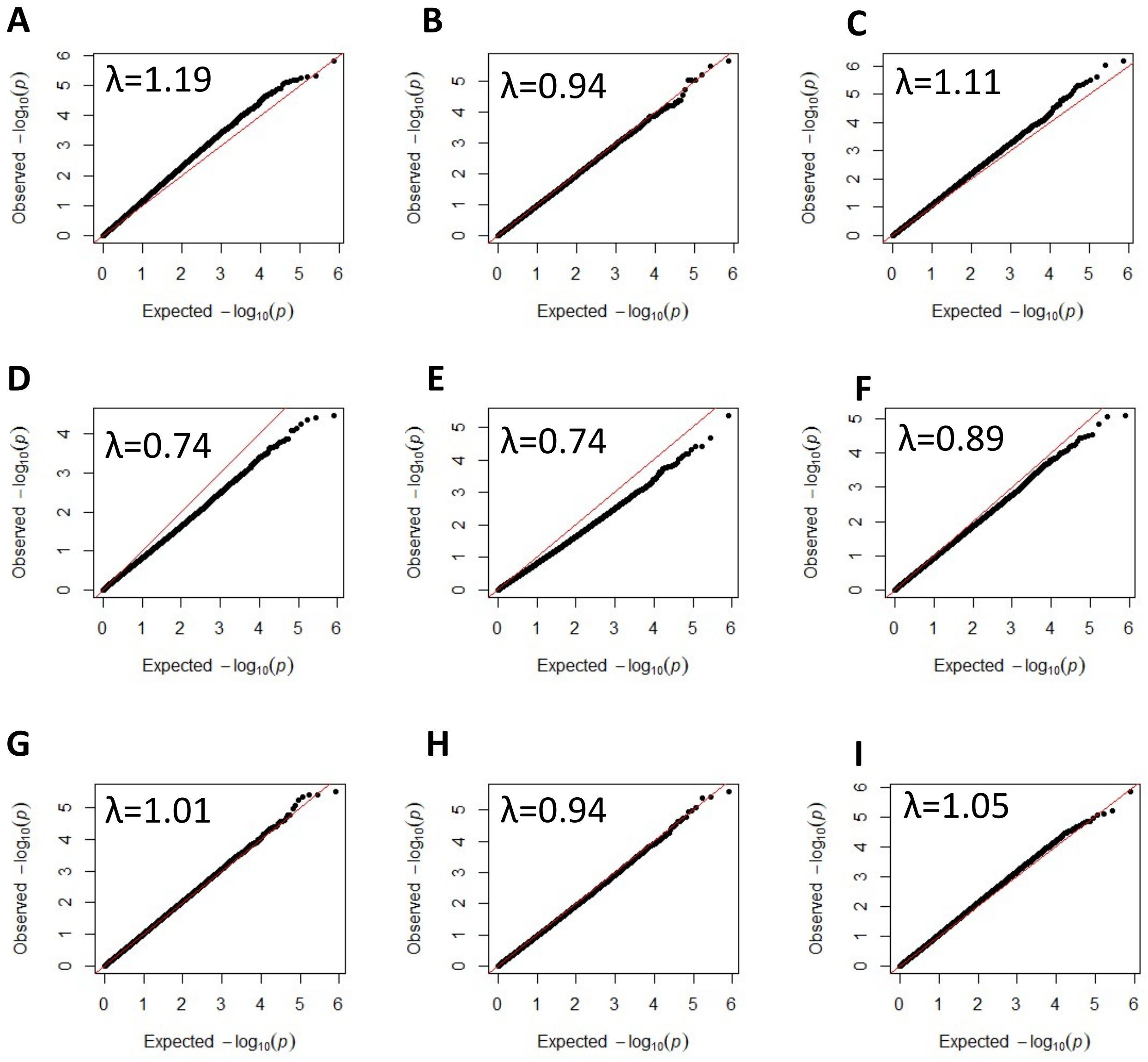
A
λ=1.19
B
λ=0.94
C
λ=1.11
D
λ=0.74
E
λ=0.74
F
λ=0.89
G
λ=1.01
H
λ=0.94
I
λ=1.05
Observed $-\log_{10}(p)$
Expected $-\log_{10}(p)$

**Figure SF1**. Sensitivity analyses: QQ plots showing the observed vs expected probabilities per CpG site from the meta-analysis of epigenome-wide association studies of cognitive skills in childhood and DNA methylation in cord blood. Each panel includes a λ index of genomic inflation. A-C) overall, verbal and non-verbal, in order, analysed as in main models and further adjustment for paternal education; D-F) overall, verbal and non-verbal, in order, adjusted as in main models and with maternal IQ instead of maternal education; G-I) overall, verbal and non-verbal, in order, as in main models with further adjustment for principal components from child's genetic data.

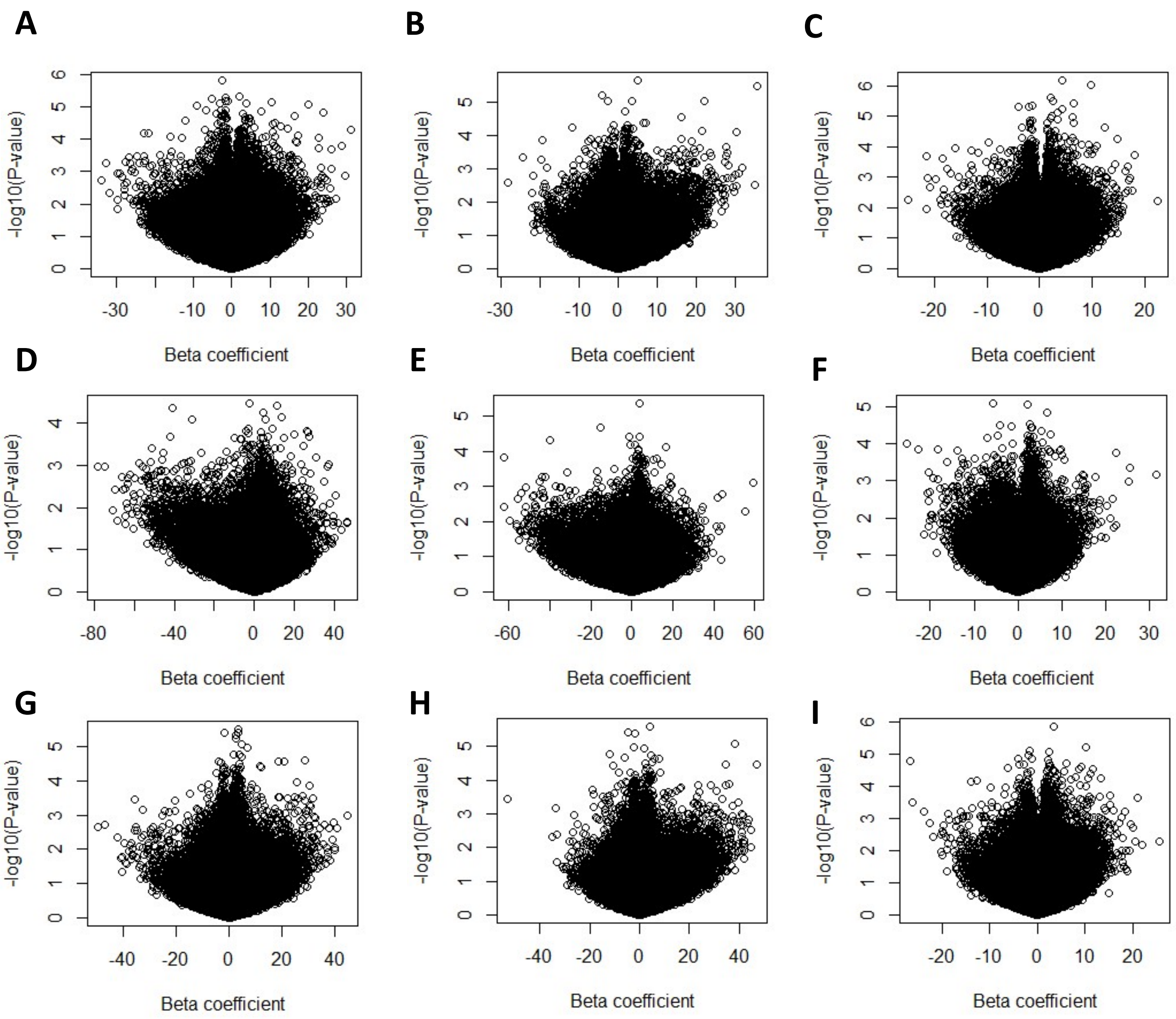
A
-log10(P-value)
Beta coefficient
B
-log10(P-value)
Beta coefficient
C
-log10(P-value)
Beta coefficient
D
-log10(P-value)
Beta coefficient
E
-log10(P-value)
Beta coefficient
F
-log10(P-value)
Beta coefficient
G
-log10(P-value)
Beta coefficient
H
-log10(P-value)
Beta coefficient
I
-log10(P-value)
Beta coefficient

**Figure SF2**. Sensitivity analyses: Volcano plots showing the effect sizes and probability values per CpG site from the meta-analysis of epigenome-wide association studies of cognitive skills in childhood and DNA methylation in cord blood. A-C) overall, verbal and non-verbal, in order, analysed as in main models and further adjustment for paternal education; D-F) overall, verbal and non-verbal, in order, adjusted as in main models and with maternal IQ instead of maternal education; G-I) overall, verbal and non-verbal, in order, as in main models with further adjustment for principal components from child's genetic data.

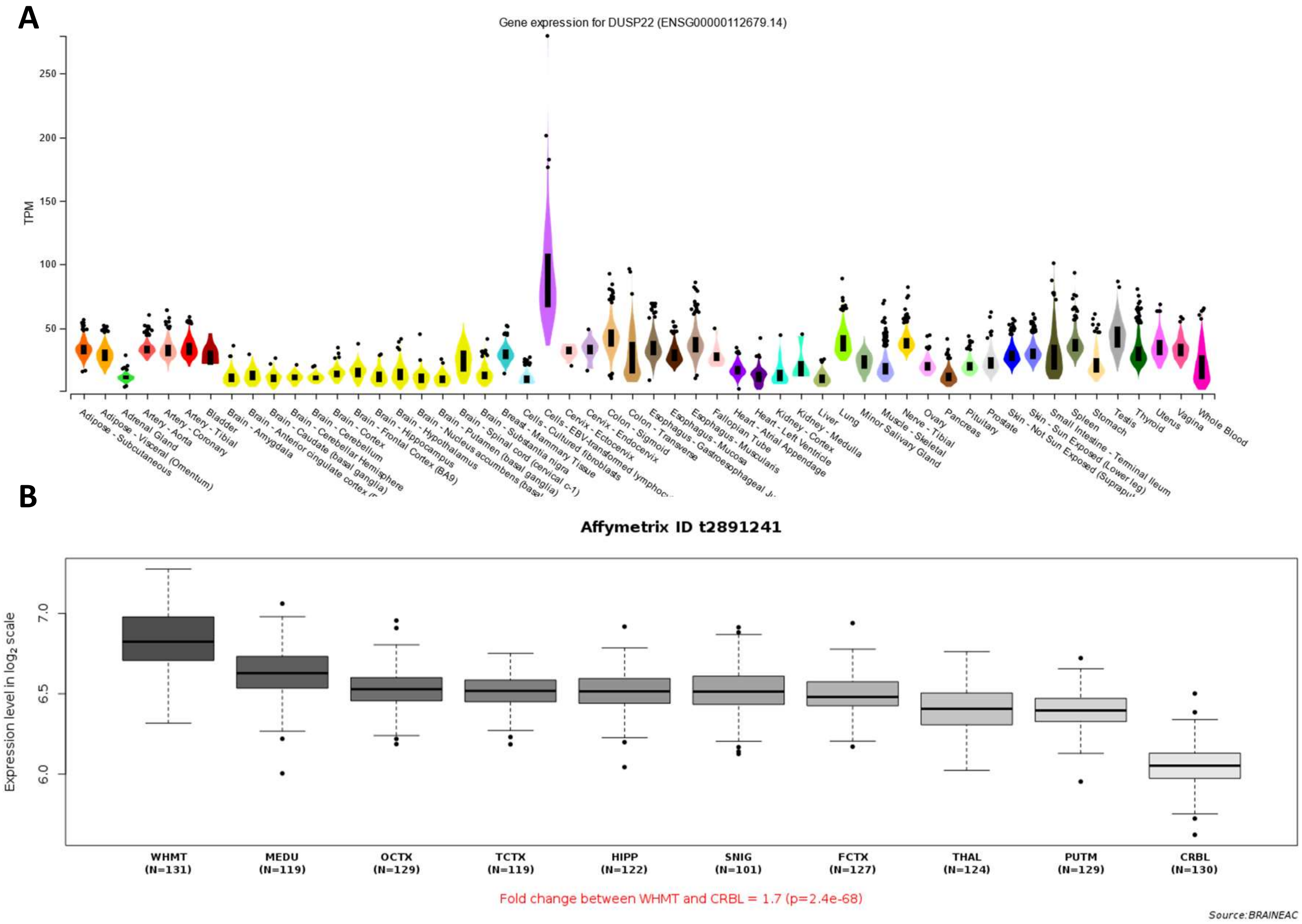
A
Gene expression for DUSP22 (ENSG00000112679.14)
TPM
250
200
150
100
50
Adipose - Subcutaneous
Adipose - Visceral (Omentum)
Adrenal Gland
Artery - Aorta
Artery - Coronary
Artery - Tibial
Bladder
Brain - Amygdala
Brain - Caudate (basal ganglia)
Brain - Cerebellar Hemisphere
Brain - Cerebellum
Brain - Cortex
Brain - Frontal Cortex (BA9)
Brain - Hippocampus
Brain - Hypothalamus
Brain - Putamen (basal ganglia)
Brain - Spinal cord (cervical c-1)
Brain - Substantia nigra
Breast - Mammary Tissue
Cells - Cultured fibroblasts
Cervix - Ectocervix
Cervix - Endocervix
Colon - Sigmoid
Colon - Transverse
Esophagus - Mucosa
Esophagus - Muscularis
Fallopian Tube
Heart - Atrial Appendage
Heart - Left Ventricle
Kidney - Cortex
Kidney - Medulla
Liver
Lung
Minor Salivary Gland
Muscle - Skeletal
Nerve - Tibial
Ovary
Pancreas
Pituitary
Prostate
Skin - Sun Exposed (Lower leg)
Small Intestine - Terminal Ileum
Spleen
Stomach
Testis
Thyroid
Uterus
Vagina
Whole Blood
B
Affymetrix ID t2891241
Expression level in log2 scale
7.0
6.5
6.0
WHMT (N=131)
MEDU (N=119)
OCTX (N=129)
TCTX (N=119)
HIPP (N=122)
SNIG (N=101)
FCTX (N=127)
THAL (N=124)
PUTM (N=129)
CRBL (N=130)
Fold change between WHMT and CRBL = 1.7 (p=2.4e-68)
Source:BRAINEAC

**Figure SF3**. *DUSP22* mRNA expression A) across tissues (source: GTEx); B) across brain areas (source: Braineac).

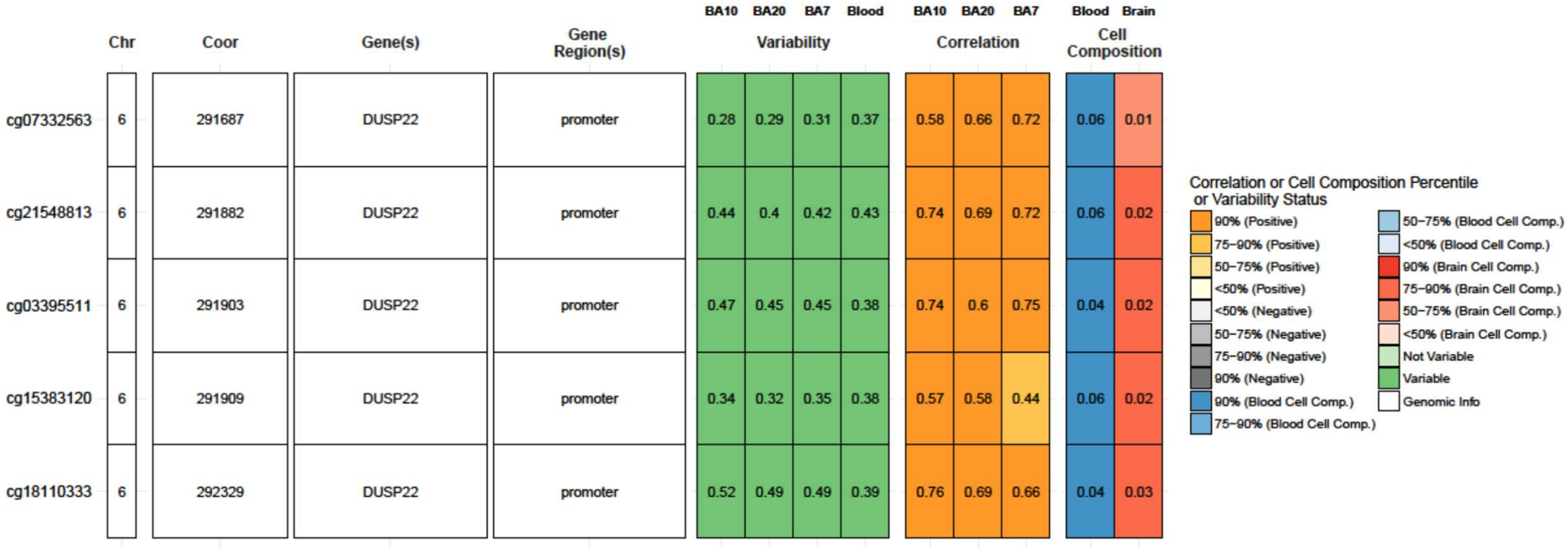


**Figure SF4**. *DUSP22* DNA methylation across brain areas (BA10, BA20, BA7) and peripheral blood (source:BECon).
